# Assessing differential cell composition in single-cell studies using voomCLR

**DOI:** 10.1101/2024.09.12.612645

**Authors:** Alemu Takele Assefa, Bie Verbist, Koen Van den Berge

## Abstract

In single-cell studies, a common question is whether there is a change in cell composition between conditions. While ideally, one needs absolute cell counts (number of cells per volumetric unit in a sample) to address these questions, current experimentation typically obtains cell counts that only carry relative information. It is therefore crucial to account for the compositional nature of cell count data in the statistical analysis. While recently developed methods address compositionality using compositional transformations together with a bias correction, they do not account for the uncertainty involved in estimation of the bias term, nor do they accommodate the mean-variance structure of the counts. Here, we introduce a statistical method, `voomCLR`, for assessing differences in cell composition between conditions incorporating both uncertainty on the bias term as well as acknowledging the mean-variance structure of the transformed data, by leveraging developments from the differential gene expression literature. We demonstrate the performances of `voomCLR`, illustrate the benefit of all components and compare the methodology to the state-of-the-art on simulated and real single-cell gene expression datasets.

## 1 Introduction

High-dimensional biological single-cell datasets, where multiple features are measured per single cell, are ubiquitous in modern biology, e.g., flow cytometry and single-cell RNA-sequencing (scRNA-seq) experiments. Usually, the features measured at cell-level (e.g., gene expression or chromatin accessibility) are used to categorize cells into cell types and/or cell states [Amezquita et al., 2020]. Conditional on such a grouping of cells, a common task concerns the estimation of cell population composition and contrasting cell population abundances between (any number of) conditions; that is, a difference in the cell type/state composition [Frishberg et al., 2019, Bues et al., 2022, Phipson et al., 2022].

Ideally, one would like to observe the absolute counts for each cell population (e.g., number of cells for each population per unit volume of blood in a patient). However, the observed data from a sequencing experiment only capture relative abundance information as the number of cells observed for a particular sample does not reflect the total number of cells in the sample [Quinn et al., 2018] but is rather constrained to an arbitrary sum. The field of compositional data analysis is concerned with performing inference on the unobserved absolute cell counts using the observed data which only contains relative information [Aitchison, 1982].

Many methodologies are described in the literature to analyze changes in cell composition. First, several approaches exist for unsupervised differential abundance analysis, which are unsupervised in the sense that no a priori known grouping of the cells is provided and instead rely on approaches like k-nearest neighbor graphs to categorize cells [Dann et al., 2022, Zhao et al., 2021, Burkhardt et al., 2021]. Here, we will instead focus on the scenario where a biologically relevant cell grouping has been provided and is followed by a differential abundance analysis. In what follows, we refer to a relevant grouping of cells as a ‘cell type’ or ‘cell population’, but note this is equally applicable for any other biologically relevant disjoint cell grouping. A common though naive approach of differential abundance analysis fits a negative binomial generalized linear model (NB-GLM) for each cell type, using the cell type counts as a response [Maity and Teschendorff, 2023, Domingo et al., 2023]. By incorporating the total number of cells per sample as offset, one models the relative abundance of each cell type. While intuitive, this approach ignores the compositional nature of the data, leading to inflated false positive rates [Zhou et al., 2022]. Indeed, compositional cell population count data are negatively correlated between populations, implying that an increase in abundance of one population will lead to a decrease of other populations, and ignoring these effects leads to an increased risk of identifying false positives. An improvement to this approach is to normalize the total cell count of each sample, and use the normalized total as offset. This is often done by using tools originally developed for bulk RNA-sequencing differential expression analysis [Weber et al., 2019], like `edgeR` and `DESeq2` [Robinson et al., 2010, Love et al., 2014]. Other methods effectively take into account the compositional nature of the data by using appropriate distributional assumptions. `DCATS` uses a Beta-Binomial regression framework for each cell type to infer differences in cell composition, where the dispersion parameter is estimated jointly across all cell types [Lin et al., 2023]. `sccomp` similarly relies on a Beta-Binomial model and allows for testing differential composition and variability between groups [Mangiola et al., 2023]. `scCODA` adopts a Bayesian multivariate regression framework based on the Dirichlet-Multinomial distribution, a generalization of the Beta-Binomial distribution. Another class of methods instead transform the cell type counts; `propeller` adopts a logit or arscin square root transformation and uses linear models post-transformation [Phipson et al., 2022]. Alternatively, compositional transformations, a class of transformations originally proposed by John Aitchision [Aitchison, 1982, 1986], are often used. They aim at transforming the data out of the simplex space, and mapping them into the real space, making measures like Euclidean distances and least squares meaningful [Quinn et al., 2018]. `LinDA` adopts a compositional transformation called the centered log-ratio (CLR) and also uses linear models downstream [Zhou et al., 2022]. The authors then show that the estimated effect sizes are biased with respect to the effect sizes one would obtain based on the absolute abundances, and propose a bias correction approach based on the mode of the effect size across cell types. Inference then occurs on the bias-corrected effect size, relying on the standard error pre-correction. While the approach was developed for differential abundance testing in microbiome data, we here show that it works very well for assessing changes in cell type composition, too.

In this paper, we compose a method for differential abundance analysis through leveraging advances of the recent literature and filling missing methodological gaps. We follow the rationale of the `LinDA` methodology by using a CLR transformation, fitting linear models, and adopting bias correction on the effect sizes. We extend this approach in two major directions. First, we show that the cell type counts are still heteroscedastic post-transformation, and account for the counts’ mean-variance structure using heteroscedasticity weights by building on the `limma-voom` framework. This has the additional advantage that we can adopt their empirical Bayes approach for shrinking linear model residual variances. Second, we account for the uncertainty involved in estimating the bias correction term by adopting a bootstrapping approach. We first discuss why these extensions are necessary in Sections 2.1 and 2.2, and then evaluate our approach, called `voomCLR`, alongside the state-of-the-art in both simulation studies (Section 2.3) and real data analyses (Sections 2.4 and 2.5).

## 2 Results

### 2.1 Compositional transformations lead to biased parameter estimates

The information content in compositional data is in the relative abundance of the features being measured [Aitchison, 1982]. It is, for example, insensible to analyze the cell counts of one particular cell type, while ignoring all other cell types, as the counts for the individual cell type contain no relevant information for analyzing compositional differences. This has inspired a range of compositional transformations that transform compositional data out of the simplex space, often through calculating log-ratios of the counts, after which standard multivariate techniques may be used for downstream analysis [Aitchison, 1982]. The denominator of the log ratio is usually either (i) the count of another feature (i.e., the count of another cell type) from the same sample, in which case the transformation is called the additive log-ratio (ALR); (ii) the geometric mean of all counts from that sample, referred to as the centered log-ratio (CLR) transformation. Various extensions to these transformations have been developed more recently, e.g., Silverman et al. [2017].

Letting *Y*_*ip*_ denote the observed cell type count of population *p* in sample *i*, these transformations may be written as

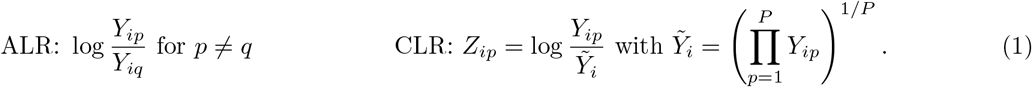

Let *X*_*ip*_ denote the (unobserved) absolute abundance of each population *p* in sample *i*, e.g., the number of T-cells in a person’s blood per volumetric unit. In the context of differential abundance (DA) analysis we are interested in testing the null hypothesis that the log-fold-change on the expected absolute abundances is equal to zero. However, when using the (observed) CLR-transformed counts *Z*_*ip*_ as a response variable in a linear model it has been shown that the obtained log-fold-changes are biased with respect to the log-fold-changes on the absolute counts *X*_*ip*_ [Zhou et al., 2022]. We indeed confirm the bias using a basic simulation study based on the Multinomial distribution (Figure 1a). In the microbiome literature, Zhou et al. [2022] propose a bias correction which we also find to alleviate the bias in our context (Figure 1a). Assuming that the majority of populations are not DA, the correction amounts to subtracting the mode of the regression coefficients across populations, a computationally tractable approach that we will be leveraging in subsequent sections. Following bias correction, the LinDA method from Zhou et al. [2022] performs inference on the bias-corrected effect sizes, assuming that the bias term was known rather than estimated. They argue that as the number of samples and the number of features tend to infinity, the uncertainty on the bias term becomes negligible as compared to the uncertainty on the (uncorrected) effect size. However, in the context of modeling cell type composition this is an unrealistic assumption as the number of cell types (features) is usually limited. Figure 1b shows that the variance on the bias correction can constitute a substantial part of the total variance (see Methods for its formulation). We therefore approximate the variance of the bias-corrected parameter by developing a bootstrap procedure to account for additional uncertainty involved in estimating the bias correction term, and incorporate it in downstream statistical inference. As can be seen in our simulation studies, this aids substantially in controlling the number of false positive results.

**Figure 1:**
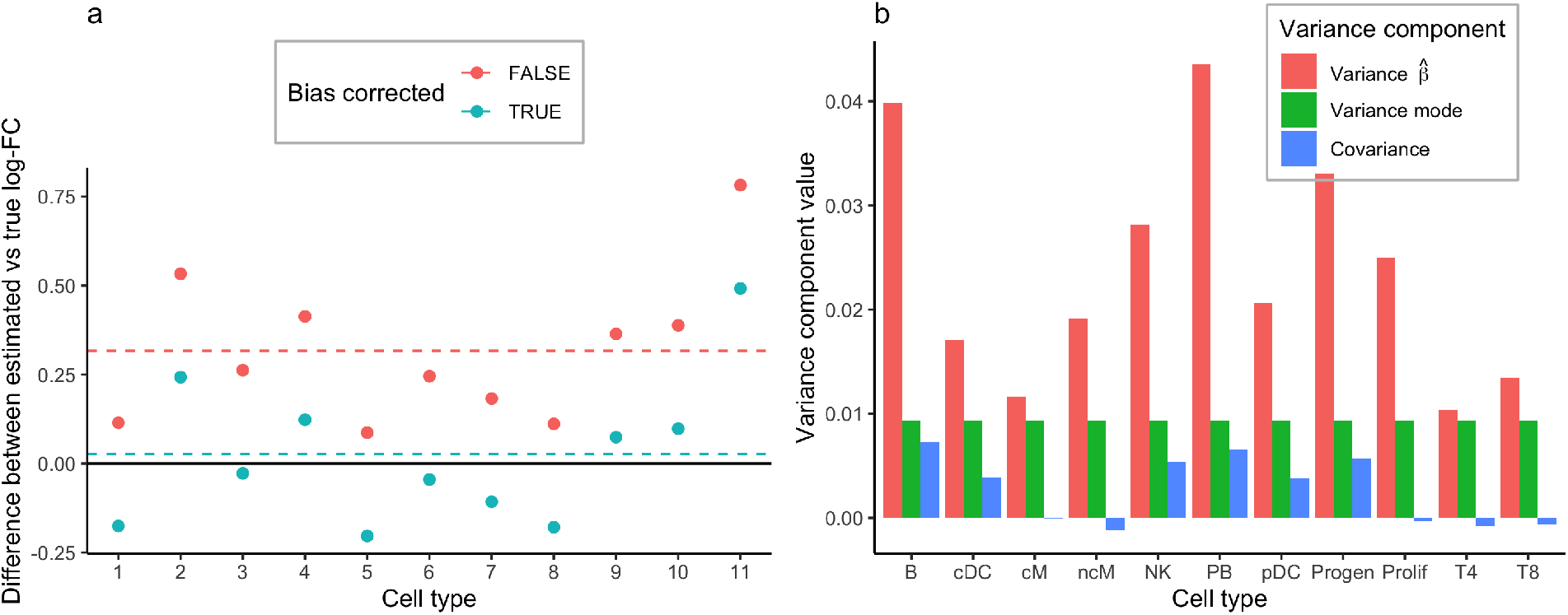
Bias correction and associated uncertainty. **(a)** Effect sizes between two groups using our parametric Dirichlet-Multinomial simulation framework. The data points represent difference of estimated minus true effect sizes (y-axis) for each cell type (x-axis). The red data points are from a `voomCLR` analysis without applying bias correction, and can be seen to be biased on average (red dashed line). The green data points are from a `voomCLR` analysis where bias correction was applied and are unbiased on average (green dashed line). **(b)** Variance components for the bias corrected effect sizes. The red bar represents the estimated variance on the estimated effect size as obtained from the weighted linear model in `voomCLR`. The green bar is the estimated variance on the bias correction term, which is the same for all cell types. The blue bar represents the estimated covariance between the two. The latter two values were estimated using our parametric bootstrap procedure.

### 2.2 Compositional transformations do not stabilize variances

The transformed counts *Z*_*ip*_ may be used downstream as the response variable in a linear model for statistical inference [Zhou et al., 2022, Hu and Satten, 2023]. However, since the untransformed data constitute counts, they have a mean-variance relationship, which is not taken into account when using a vanilla linear model on the transformed data. We explore the mean-variance relationship using both a null simulation study assuming a Multinomial distribution for *Y*_*ip*_ as well as real data. Figure 2a shows that, for the Multinomial simulation, the mean-variance relationship for the cell population counts across samples can be approximated by a Poisson law (see Methods). After CLR transformation, the variance is not independent of the mean, and indeed is a decreasing function of the mean CLR-value (Figure 2b). We see similar patterns using our parametric Dirichlet-Multinomial simulation framework (Figure 2c-d), as well as using real scRNA-seq data (Figure 2e-f). Using the first-order delta method [Dorfman, 1938, Doob, 1935] (see Methods) one can approximate the variance of the transformed data to theoretically show it is still a function of the mean, i.e., the compositional transformations are not variance-stabilizing.

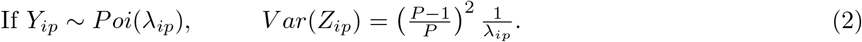

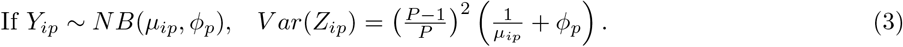

**Figure 2:**
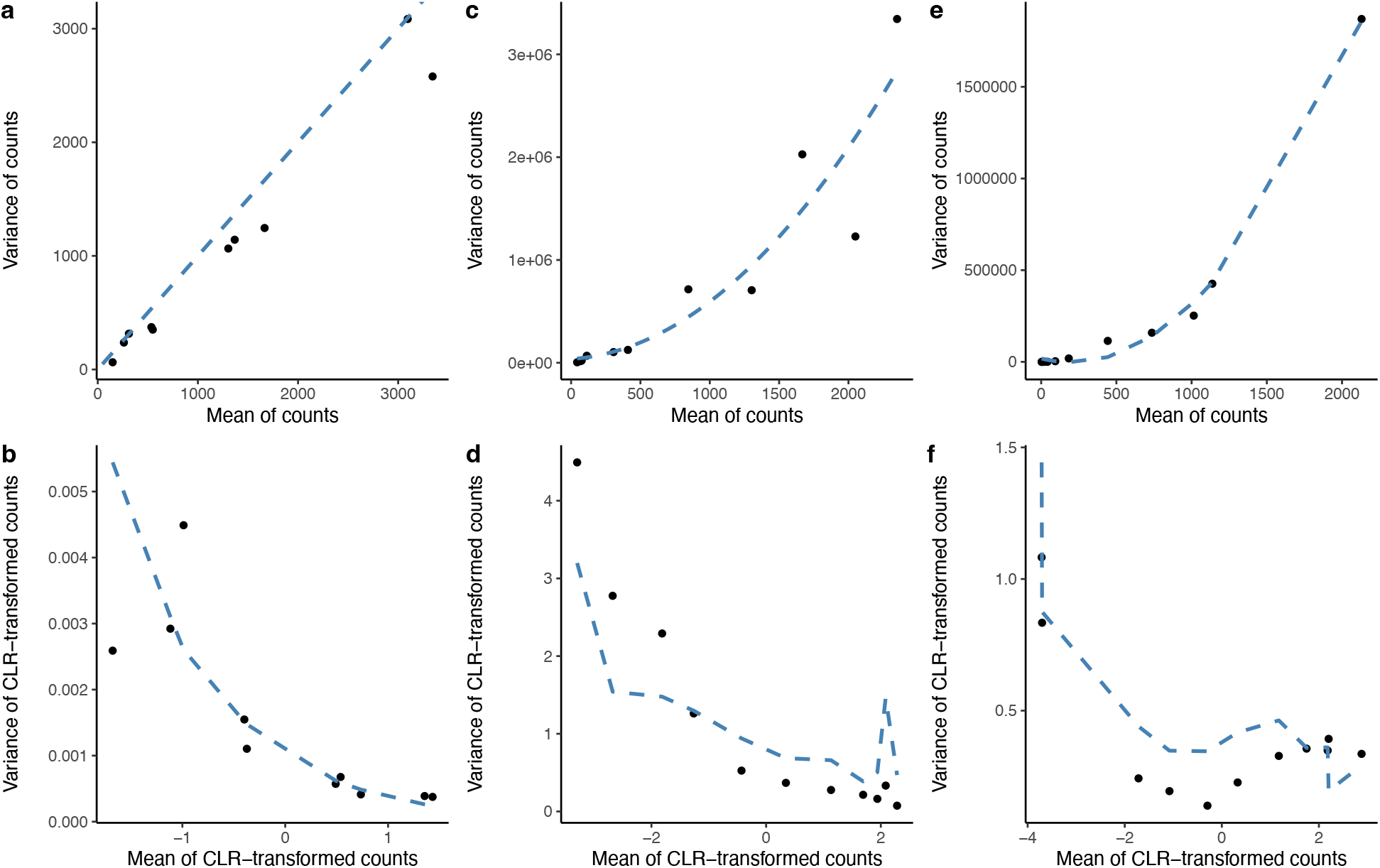
Heteroscedasticity for simulated and real data. **(a)-(b)** Simulation based on Multinomial distribution: Panel (a) shows the empirical mean-variance relationship of the counts for each cell type, with the blue line being the identity line. Panel (b) shows the empirical mean-variance relationship of the CLR-transformed cell type counts, with the blue line being the analytically calculated variance using the Delta Method based on the Poisson distribution. **(c)-(d)** Simulation based on a Dirichlet-Multinomial distribution: Panel (c) shows the empirical mean-variance relationship of the counts for each cell type, with the blue line being the estimated mean-variance trend. Panel (d) shows the empirical mean-variance relationship of the CLR-transformed cell type counts, with the blue line being the analytically calculated variance using the Delta Method based on the negative binomial distribution. **(e)-(f)** Lupus dataset, upon selecting the healthy patients from processing cohort 1: Panel (e) shows the empirical mean-variance relationship of the counts for each cell type, with the blue line being the estimated mean-variance trend. Panel (f) shows the empirical mean-variance relationship of the CLR-transformed cell type counts, with the blue line being the analytically calculated variance using the Delta Method based on the negative binomial distribution. Note that the trend in panels (d) and (f) are not smooth since they rely on a cell type specific dispersion estimate.

To address the heteroscedasticity of *Z*_*ip*_, our approach is inspired by the work from Law et al. [2014] who address the issue for transformed RNA-seq count data by estimating an empirical mean-variance trend across all genes, and leverage this trend to incorporate the mean-variance relationship through observation-level weighting in a linear model. Here, we adapt their strategy to accommodate the centered log-ratio transformation, and empirically estimating the mean-variance trend across cell populations (instead of genes). By further leveraging the software implementation from Law et al. [2014], we also gain by adopting the empirical Bayes shrinkage of linear model residual variances, across cell types. This helps especially in small sample size settings. However, in contrast to RNA-seq data where one observes thousands of genes, some datasets may only consist of a limited number of cell populations, rendering the empirical estimation of the mean-variance trend uncertain. We therefore also provide the option to derive the weights analytically, using the Delta method (see Methods).

### 2.3 Simulation studies

In this section, we will first evaluate the performance of `voomCLR` with respect to different options of accounting for heteroscedasticity and uncertainty of the bias correction. Next, we compare `voomCLR` to popular and state-of-the-art statistical methodology to assess differences in cell type composition. The evaluation centers around two key characteristics. First, the false discovery rate (FDR) quantifies the expected fraction of incorrectly identified differentially abundant cell populations among all discoveries. Second, the sensitivity level (also known as the true positive rate or TPR) assesses each method’s ability to detect truly differentially abundant cell populations. Both of these characteristics are computed at a specified nominal FDR level. An ideal yet realistic method exhibits an FDR close to or below the nominal level, with a TPR as high as possible, ideally approaching 1. We employed two approaches to simulate realistic cell count data: (1) a parametric simulation in which cell counts are generated from a Dirichlet-Multinomial distribution; (2) a non-parametric framework in which signal is introduced in a real dataset. We limited the simulations to a two-group comparison of independent replicates. The details of these simulation methods can be found in the Methods section. Supplementary Figures S1 -S5 visualize the comparison between a real and a representative simulated dataset, showing simulated data captures essential characteristics of real data.

#### 2.3.1 Performance of voomCLR

Figure 3 shows the performance of `voomCLR` across 9 different configurations of the method. These include three options to accommodate the estimation uncertainty of the bias correction factor (none, non-parametric bootstrap or parametric bootstrap) combined with three approaches to estimate the observation-level heteroscedasticity weights (empirical weights or analytically calculated weights based on either a Poisson or negative binomial distribution). In a general view of the results, most of the `voomCLR` settings demonstrated reasonable control of FDR and achieved a high TPR for testing differential abundance. In particular, the results indicate that accounting for the uncertainty of the bias correction factor using the bootstrap improves the performance of `voomCLR` to control the FDR level near or below the nominal level. The FDR level from the non-parametric bootstrap procedure was lower than that of the parametric procedure, albeit accompanied with a somewhat lower TPR. Furthermore, `voomCLR` with analytical observation-level weights based on the negative binomial distribution outperforms the configuration using the Poisson distribution. The latter often fails to maintain FDR control and exhibits a lower TPR in general. The performance difference among the different configurations of `voomCLR` is more pronounced for small and moderate sample sizes. Supplementary Figure S7 further compares the performance of these configurations at varying levels of simulated biological variability between samples (low, medium, and high) and three different sample sizes, based on the parametric simulation. As one could expect, the results primarily indicate that larger variability negatively impacts methods’ TPR, which however improves with increased sample size. Notably, even at at high biological variability, `voomCLR` configurations using bootstrap procedures to account for bias correction uncertainty demonstrated better FDR control.

**Figure 3:**
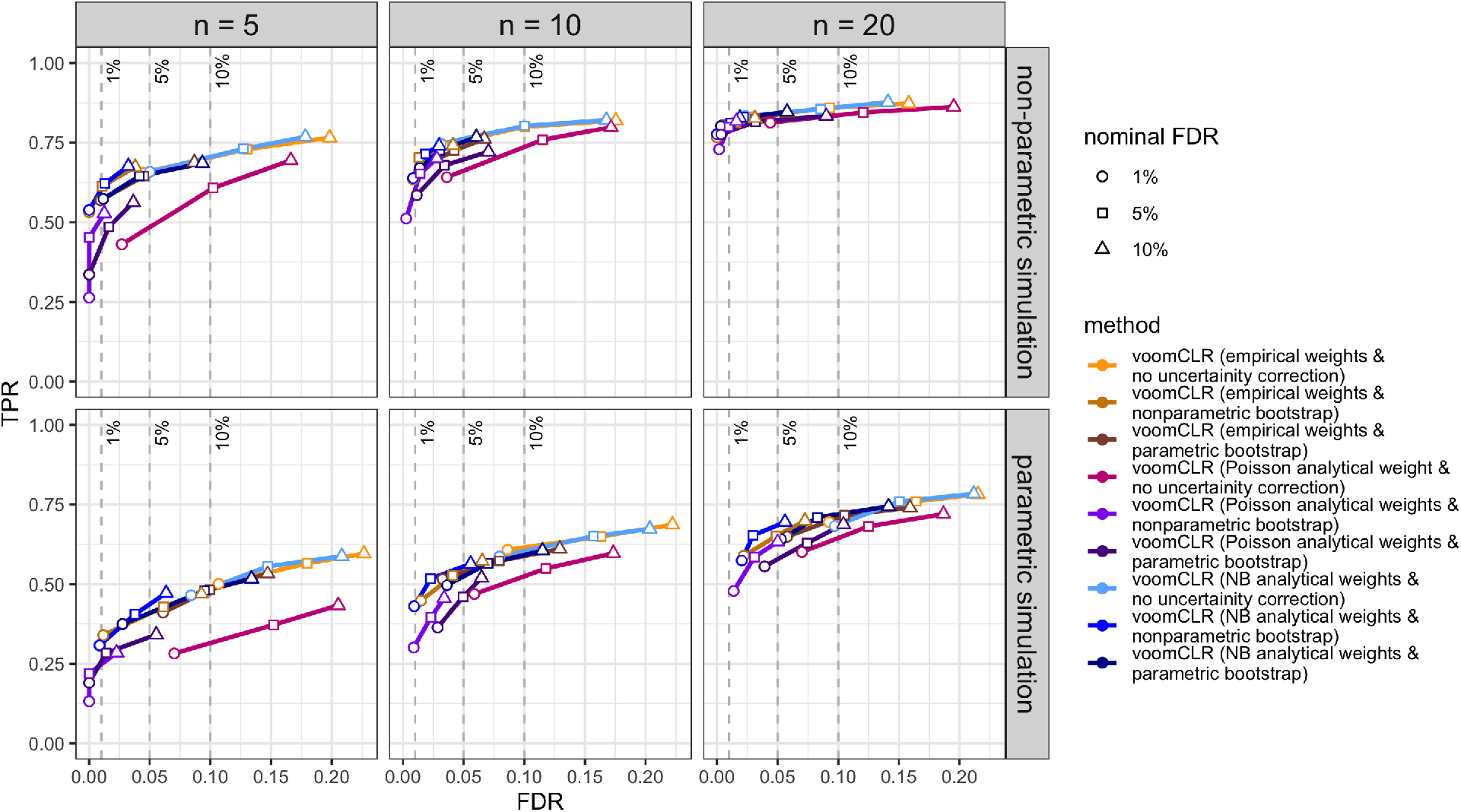
FDR-TPR performance curves of nine different voomCLR configurations in two simulation paradigms. The curves indicate the performance of the methods for testing differential abundance in simulated compositional cell count data. Performance metrics (FDR and TPR) are calculated at 1%, 5% and 10% nominal FDR levels for each simulation setting. Two simulation strategies are used: (1) non-parametric simulation based on recycling real cell abundance data (using the lupus dataset from Perez et al. [2022]), and (2) parametric compositional cell count data simulation using a Dirichlet-Multinomial distribution. Reported metrics are averages from 250 simulation runs for each simulation scenario. Each simulated dataset consists of two independent groups of samples for 11 cell populations. Data is simulated with a sample size of 5, 10 and 20 per group and with a medium level of variability between samples (Dirichlet parameters scaling factor, *γ* = 1). NB = negative binomial.

Moreover, the simulation result in Supplementary Figure S11 further compares the performance of `voom-CLR` configurations for an increasing number of cell types. The results indicate that `voomCLR` with empirical observational weights tend to lose control of FDR when the number of cell types P is less than 10, probably owing to the high degree of uncertainty in the empirical mean-variance trend estimation. However, the negative binomial-based analytical weights effectively control the FDR while maintaining a high TPR. For medium to high numbers of cell types, the choice between empirical and negative binomial-based analytical weights does not have a significant impact. On the other hand, Poisson-based analytical weights generally underperformed concerning TPR level in most of the simulation settings.

Generally, the non-parametric bootstrap procedure is conservative in terms of FDR control and therefore comes with an associated cost of reduced sensitivity. On the other hand, the parametric bootstrap is moderately liberal and is associated with higher sensitivity. The simulation study indicated that given `voomCLR` is applied with a parametric bootstrap procedure (for *P* ≤10), it is best to use the analytically calculated observational weights based on the negative binomial distribution for optimal performance. In addition, these results indicate that the choice between parametric and non-parametric bootstrap is more important than the choice between Poisson and negative binomial distribution for observation weights. For *P* ≤ 10, the choice between empirical or negative binomial based analytical weights has minimal impact on the performance of `voomCLR`. As a result, we recommend `voomCLR` either with parametric bootstrap and empirical observational weights (`voomCLR (empirical weights & parametric bootstrap)`) or parametric bootstrap procedure with analytical observational weights based on the negative binomial distribution (`voomCLR (NB analytical weights & parametric bootstrap)`) for differential abundance analysis. These conclusions generally apply to the results from both simulation procedures (non-parametric and parametric). However, the TPR of all methods observed in the parametric simulation study is relatively lower. One possible explanation for this difference is that the variability (between or within cell populations) in the parametric simulated data is higher than that in the non-parametric simulation (Supplementary Figures S2 -S5).

In order to also evaluate the impact of adopting bias correction and empirical Bayes shrinkage of the residual variances, we also evaluated the performance of `voomCLR` with and without these features. The results shown in Supplementary Figure S9 indicate the heteroscedasticity correction improved the TPR level of `voomCLR` while the empirical Bayes procedures improved the FDR control. These procedures improve the FDR control of `voomCLR` only when applied together with the bias correction step. Nevertheless, for a relatively large sample size (*n* ≥ 20), none of these procedures contribute significantly to the performance of FDR control and TPR except for the bias correction step.

Based on the performance evaluation discussed above, `voomCLR (NB analytical weights & parametric bootstrap)` is chosen as a default method in the subsequent performance evaluations presented in the paper; we will use the shorter name `voomCLR` to refer to this configuration.

#### 2.3.2 Performance of voomCLR compared to other methods

The simulation study further allows evaluating the performance of `voomCLR` compared to other popular methods for differential abundance analysis. The methods included in the evaluations can be categorized into three groups. First, methods that apply linear models on CLR-transformed cell counts. Under this category, we have `voomCLR`, `LinDA` [Zhou et al., 2022], and as a baseline a vanilla linear model using the CLR-transformed counts as response (`LM CLR`) – fits the linear model and makes inference based on the estimated effect sizes with neither compositional bias nor heteroscedasticity correction. In the second category, we have methods developed for differential expression analysis of digital gene expression data. The methods under this category are `edgeR` [Robinson et al., 2010], `DESeq2` [Love et al., 2014] and `limma-voom` [Law et al., 2014]. `edgeR` and `DESeq2` employ a negative binomial generalized linear model (NB-GLM) framework for testing differential abundance. `limma-voom` fits linear models on log-transformed counts per millions (CPM). It shares various techniques, such as normalization of total cell counts and moderated tests, with `edgeR` and `DESeq2`. As a baseline for the second category, we have included ordinary negative binomial GLMs (`NB_GLM (TMB)`) which use the logarithm of the total cell count per sample as an offset, fitted using the `glmmmTMB` R package [Brooks et al., 2017]. The third category includes methods specifically developed for testing differential abundance in compositional cell population data often originating from single-cell RNA-seq data. These include `propeller` (with logit or arcsine transformation of cell counts) [Phipson et al., 2022] and `DCATS` [Lin et al., 2023]. A brief description of all these methods and implementation notes can be found in the Methods section.

The results presented in Figure 4 compare the 10 methods discussed above based on a simulation setting with 11 cell populations (*P* = 11) at 3 different simple sizes (*n* = 5, 10 and 20 in each group), similar to Figure 3. Notably, two methods—`voomCLR` and `LinDA`—stand out with respect to FDR control compared to all the other methods. Recall that these methods model CLR-transformed data and incorporate bias correction for effect sizes. In contrast, the linear model applied to CLR-transformed counts without bias correction, heteroscedasticity correction, or empirical Bayes shrinkage performs poorly in terms of both FDR control and TPR. Although CLR transformation is widely used to process compositional data, our results suggest that relying solely on CLR transformation for compositional cell count data does not guarantee effective control of false positives when one tests for differential abundance.

**Figure 4:**
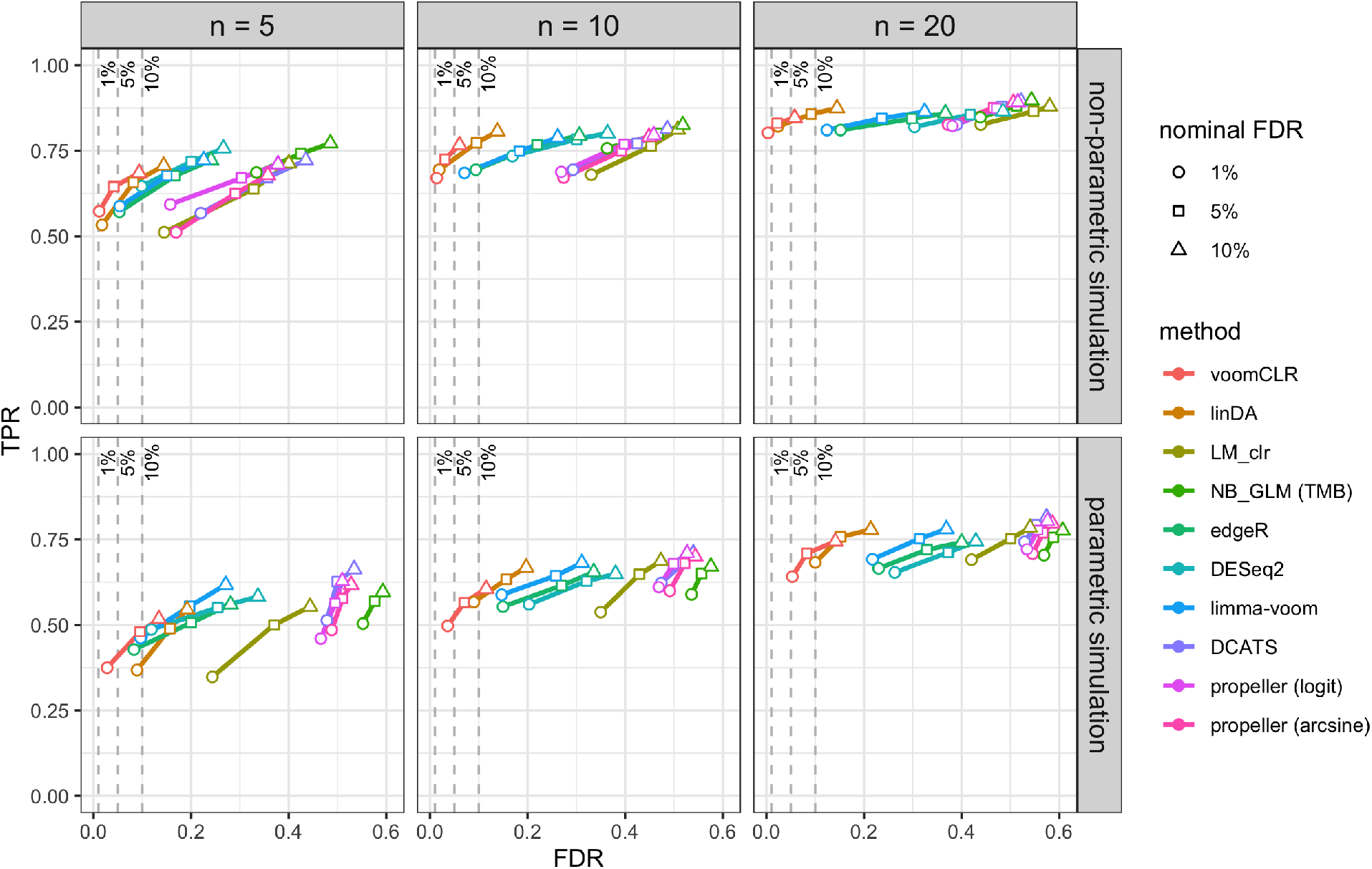
FDR-TPR performance curves of 10 methods from two simulation paradigms: The curves indicate the performance of the methods for testing differential abundance in simulated compositional cell count data. Two simulation strategies are used: (1) non-parametric simulation based on recycling real cell abundance data (using the lupus dataset from Perez et al. [2022]), and (2) parametric compositional cell count data simulation using a Dirichlet-Multinomial distribution. Performance metrics (FDR and TPR) are calculated at 1%, 5% and 10% nominal FDR levels for each simulation setting. Reported metrics are averages from 250 simulation runs for each simulation scenario. Each simulated dataset consists of two independent groups of samples for 11 cell populations. Data is simulated with a sample size of 5, 10 and 20 per group and with a medium level of variability between samples (Dirichlet parameters scaling factor, *γ* = 1).

In particular, the results in Figure 4 indicate that `voomCLR` generally outperforms `LinDA` especially when the sample size is low (*n* ≤ 10). This is particularly apparent when the variability between samples is medium to high, and holds even for 10 to 20 samples per group. (Supplementary Figure S8). In simulation scenarios with medium and large variability between samples, `voomCLR` demonstrates a significant performance advantage in controlling the FDR compared to `LinDA` even for 10 and 20 samples per group. `voomCLR` achieves this while maintaining a comparable TPR for detecting truly differentially abundant cell populations. Although both tools leverage the bias correction procedure, `voomCLR`’s superior performance can be explained by its additional incorporation of heteroscedasticity, empirical Bayes shrinkage of the residual variances and accounting for the uncertainty in the bias correction. One assumption for the bias correction procedure is that the fraction of truly differentially abundant cell populations is not large. However, an additional simulation study spanning low to high proportions of truly differentially abundant cell populations (Supplementary Figure S6) demonstrates that the performance of `voomCLR` in controlling the FDR remains reasonably well even when the fraction of truly differentially abundant cell populations is large.

Another popular and naturally intuitive approach for testing differential abundance is the negative binomial GLM framework, using the total cell counts as an offset (`NB_GLM (TMB)`). However, our simulation study shows that this approach generally leads to poor performance, with the actual FDR often exceeding 50% for a nominal 5% FDR (Figure 4 and Supplementary Figure S8). Although it achieved the highest TPR compared to all the methods evaluated in the study, the inadequate FDR control suggests that it is an ineffective procedure for handling compositional data. Methods developed for differential gene expression analysis, `edgeR`, `DESeq2` and `limma-voom` demonstrated improved performance compared to that of a vanilla NB-GLM. The improved performance of `edgeR`, `DESeq2` and `limma-voom` can be explained by the normalization procedure these three tools apply on the total cell counts that helps in dealing with compositional data and the empirical Bayes procedure for moderated testing (Supplementary Figure S10). Nevertheless, for a typical number of cell populations that is between 10 and 30, `edgeR`, `DESeq2` and `limma-voom` underperform compared to `voomCLR` in terms of FDR control whereas, in simulated data with more than 50 cell populations, these methods perform relatively well and comparable to `voomCLR` (Supplementary Figure S12). This can be explained by the fact that the compositional effect decreases when the number of cell populations increases.

Moreover, the simulation study indicates that `propeller`–a recently introduced tool designed specifically for testing differential abundance in compositional cell population data from single-cell RNA-seq data – generally did not perform well in both simulation settings (Figure 4 and Supplementary Figures S8 and S12). Specifically, the FDR level far exceeds the nominal level indicating its inadequacy for properly handling compositional effects. The choice of transformation techniques (arcsine or logit) did not significantly alter `propeller`’s performance. Similarly, `DCATS` – also developed for testing differential abundance in compositional cell population data from single-cell RNA-seq data using a beta-binomial GLM framework– ranked equally to propeller with respect to FDR control and TPR in all simulation settings (Figure 4 and Supplementary Figures S8 and S12).

Finally, we repeated the non-parametric simulation study using the healthy samples from the Breast Cell Atlas data as a baseline. The results shown in Supplementary Figure S13 further confirm the superior performance of `voomCLR` over the other methods evaluated in the study with respect to FDR control. In this particular simulation, all methods generally have lower TPR. However, from assessing the mean-variance trend, it can be seen that there is a larger biological variability in the Breast Cell Atlas data than in the Lupus data (Supplementary Figure S14) which negatively impacts the performance of methods especially at low sample sizes. This result is consistent with the observation from the parametric simulation study with high simulated biological variability (Supplementary Figure S8). The results in Supplementary Figure S13 also indicate that `LinDA` performs relatively poorly with respect to FDR control in this simulation study especially when the sample size is 5 per group. `Propeller` (with both transformation methods) on the other hand showed relatively better performance for FDR control. In summary, `voomCLR` is a robust, powerful method that, across a broad range of scenarios and simulation frameworks, dominates other methods.

### 2.4 Lupus case study

Perez et al. [2022] perform scRNA-seq on 355 human blood samples from 261 individuals, corresponding to 99 healthy controls and 162 systemic lupus erythematosus (SLE), hereafter also referred to as ‘lupus’, patients. Following the analysis from the original paper, we here focus on samples from humans with Asian and European ancestry. After quality control and initial filtering, more than than 1.2 million cells remained, which were assigned to 11 cell types [Perez et al., 2022]. We perform exploratory data exploration consistent with compositional data analysis by using a principal component analysis based on Aitchison’s distance. While the healthy samples are more tightly grouped as compared to SLE samples (Figure 5a), both groups overlap. We find no known technical variation to be associated with the variance captured in the reduced dimension plot (Supplementary Figure S15). The CLR-transformed data are shown in Figure 5b. We model the data using `voomCLR` accounting for processing cohort as a covariate, and also include an interaction between lupus status and ancestry, as in the original paper the lupus status had been assessed for samples from Asian and European ancestry separately. We account for replicate sequencing of some individuals through the `duplicateCorrelation` feature from `limma`. Given the superior performance of the parametric bootstrap in our non-parametric simulation study based on this dataset, we account for uncertainty of the bias correction using the parametric bootstrap. Statistical inference is performed on a 10% nominal false discovery rate (FDR) level, testing the lupus disease effect within Asian and European ancestries, as well as the interaction between lupus disease status and ancestry. In doing so, we confirm the finding of the original paper that, for both ancestries, the average abundance of cM and proliferating cells increases in lupus versus control samples, while additionally finding that the average abundance for non-classical monocytes increases in lupus samples versus controls; see Figure 5c for a visual summary. Cell types found to be different in average abundance only in the European ancestry were plasmacytoid dendritic cells (pDCs). For the Asian population, we find a decrease of CD4 T-cells (T4) and a decrease in CD8 T-cells (T8). Thus, validation of the original analysis on the CD4 T-cells is only confirmed for Asian ancestry, but not for European ancestry. Interestingly, the original analysis notes that the decrease in CD4 T-cell abundance is higher for Asian cases as compared to European ones, which the authors confirm using external data based on complete blood counts. When testing for an interaction effect between disease and ancestry, we find no significant cell types, however, CD4 T-cells are indeed the top ranked cell type, however, with an unadjusted p-value of 0.51 (FDR-adjusted p-value of 0.60).

**Figure 5:**
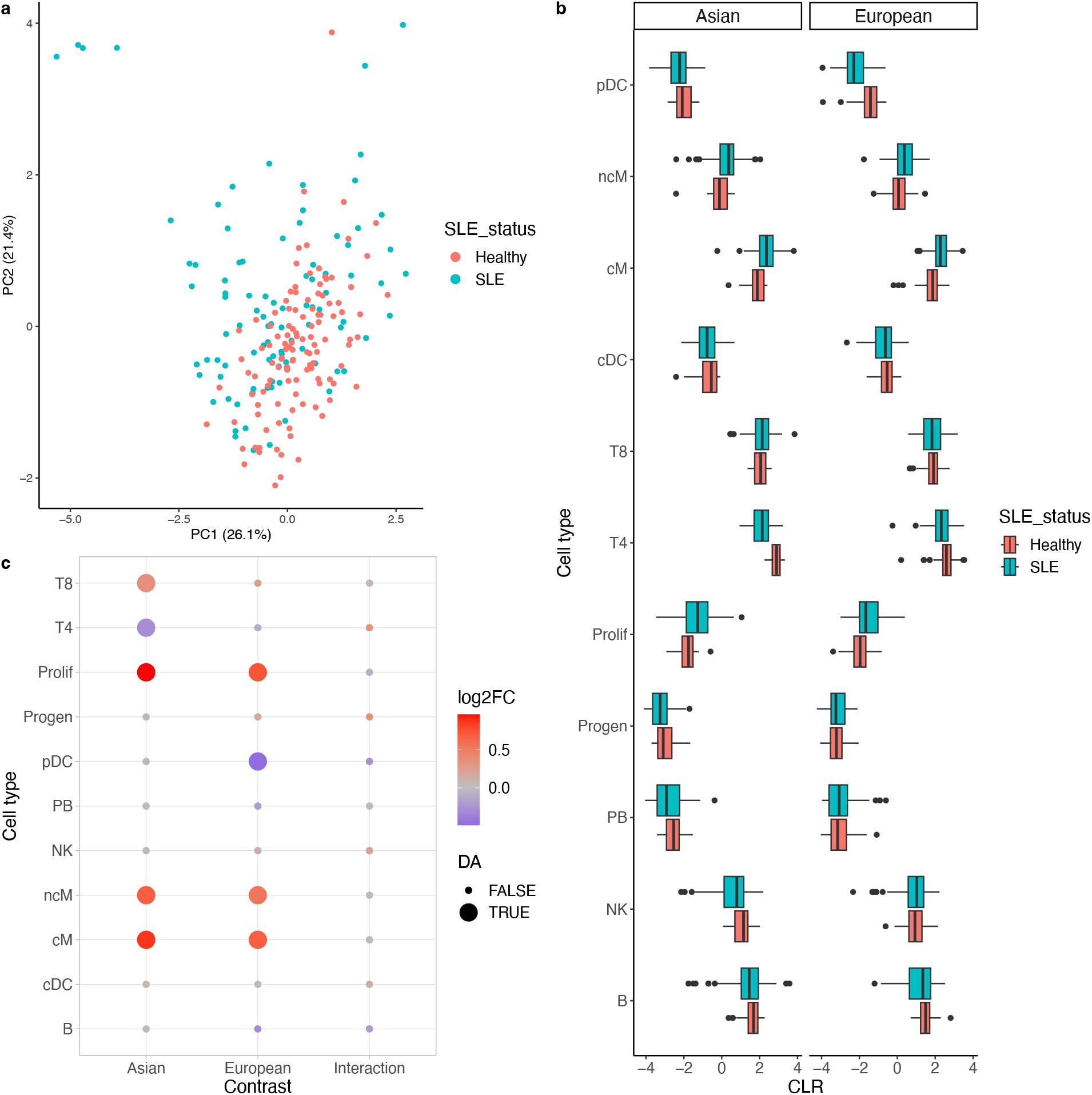
Lupus case study. **(a)** Scatterplot of first two principal components of Aitchison’s distance PCA for all samples. Each sample is colored by disease status. **(b)** Boxplots of CLR-transformed cell type abundances, for both the Asian and European ancestries. **(c)** Heatmap showing results of `voomCLR` for all cell types (y-axis). Each point is colored according to the log2 fold-change, and point size denotes whether the FDR-adjusted p-value is below the 10% level.

We note that these results are largely due to a switch in analysis methodology, rather than accounting for the processing cohorts in the statistical analysis. When not accounting for the processing cohorts, in terms of statistical significance on a 10% FDR level, the qualitative results mostly remain for the disease effect within each ancestry (Supplemental Figure S16). Only one additional finding is that B-cells decrease in European lupus patients versus the healthy population. For the interaction effect, we now do pick up the CD4 T-cells and additionally also pick up progenitor cells. These findings suggest that the CD4 T-cell interaction effect picked up in the original manuscript may partially be driven by technical effects from the processing cohorts.

We contrast our results against two alternative methods, the negative binomial GLM (NB-GLM) and LinDA (see Methods) for all three contrasts. Detailed results are shown in Supplementary Figure S17. In general, cell types that are discovered by `voomCLR` are confirmed by other methods. However, other methods often uniquely identify additional cell types as well, possibly reflecting the increased false discovery rate of those methods, which may be alleviated using our bootstrapping approach. Across all 33 evaluations (11 cell types × 3 contrasts), all methods agree on the DA calling at a 5% FDR level for 27 of them, showing major agreement. The NB-GLM has unique DA calls for 4 evaluations, while voomCLR and LinDA agree for 4 evaluations, where NB-GLM has a different result (Supplemental Figure S18).

### 2.5 Human breast cell atlas case study

A cell atlas of the human breast was recently published by Reed et al. [2024], where a cohort of healthy breast tissue samples from 55 donors was assessed using single-cell RNA-sequencing. The cohort constitutes 22 women having undergone reduction mammoplasty, 27 samples from women who carry a *BRCA1* or *BRCA2* mutation or had a family history of breast cancer that was not attributed to known risk genes, and 6 *BRCA1*-carrier women that had breast cancer in one breast and had the second breast removed to reduce risk of further tumors [Reed et al., 2024]. A total of 800, 000 single cells were measured, which could be split in three major compartments: stromal, epithelial and immune cells.

Like the original manuscript, we will investigate the association of genetic and environmental factors with cell type composition in the breast. In particular, we assess the impact of genetic risk factors, comparing high-risk *BRCA1* - and *BRCA2* -carriers with average risk donors; see Methods. These contrasts were assessed for the stromal, epithelial and immune cells separately, like in the original manuscript [Reed et al., 2024]. However, here we will focus on immune cells (Figure 6a), as a subset of these results were validated in follow-up experiments, providing a near ground truth. Indeed, enrichment of CD8- and CD4 T-cell types in high risk (HR) donors as compared to average risk (AR) donors was confirmed via immunofluorescence staining. Alongside `voomCLR`, we also compare with methods `LinDA` and `edgeR` as well as the neighborhood-based `milo` approach [Dann et al., 2022] used in the original paper.

**Figure 6:**
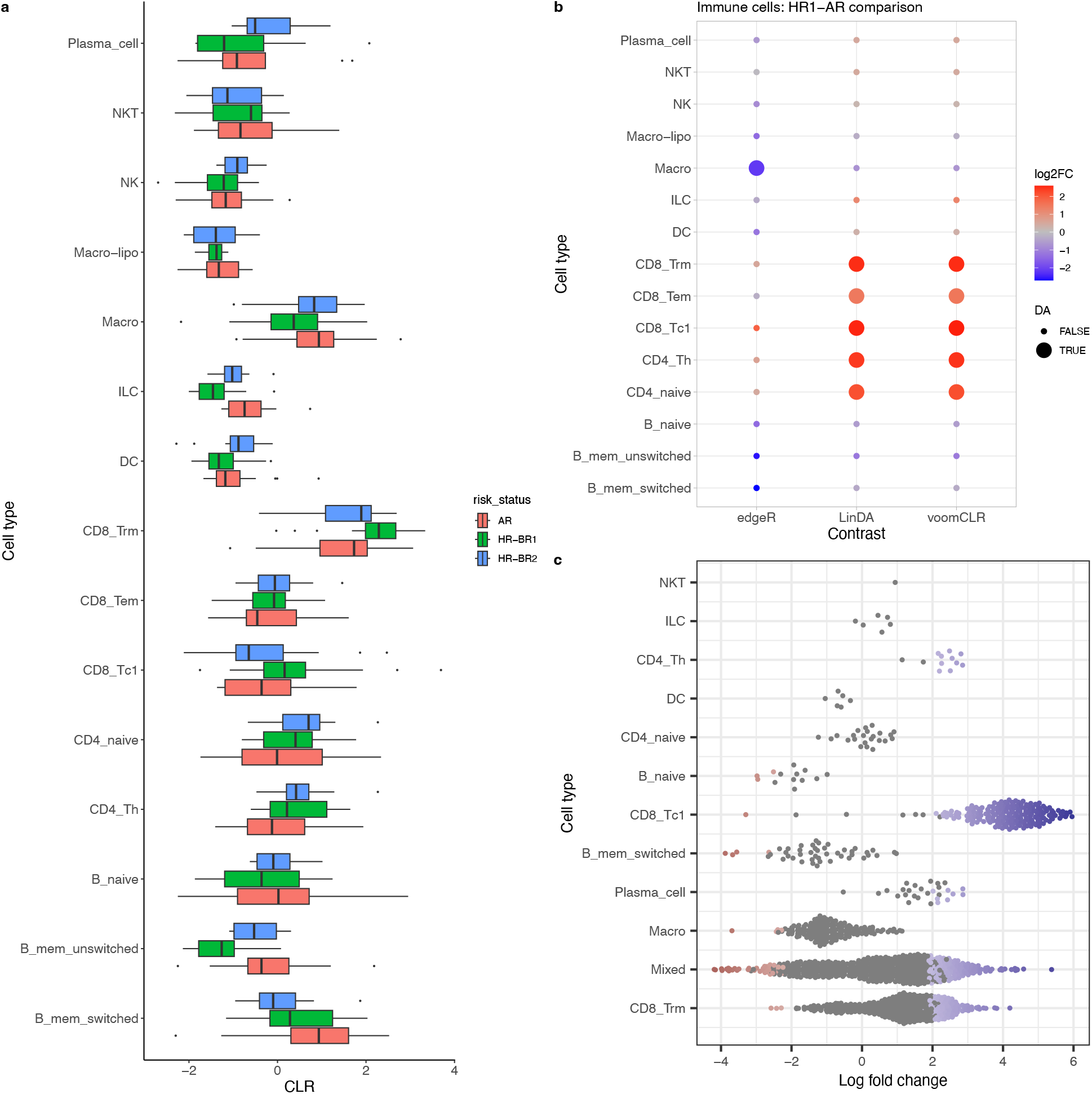
Breast Atlas case study. **(a)** Boxplots of CLR-transformed cell type abundances, for average risk (AR) donors and high risk donors with BRCA1 (HR-BR1) and BRCA2 (HR-BR2) mutations. **(b)** Heatmap showing results of `edgeR`, `LinDA` and `voomCLR` without bootstrapping, for all cell types (y-axis) in the comparison of HR-BR1 versus AR donors. Each point is colored according to the log2 fold-change (positive is higher abundance in HR-BR1 donors), and point size denotes whether the FDR-adjusted p-value is below the 5% level. **(c)** `Milo` results for all cell types, where each data point represents a neighborhood. Neighborhoods are assigned to a cell type as soon as at least 70% of its constituent cells are from that cell type. Significant neighborhoods (5% FDR level) are colored, while insignificant ones are in grey.

In this comparison, `voomCLR` and `LinDA` confirm that CD8 and CD4 T-cell types are increased in both high risk (HR) donor groups as compared to the average risk (AR) donors, while `edgeR` does not find any significant effects for these cell types (Figure 6b and Supplementary Figure S19); underscoring the importance of accounting for compositionality. For both comparisons, the bias correction term is fairly uncertain, rendering some cell types insignificant at 5% FDR level when using the parametric bootstrap in a `voomCLR` analysis (Supplementary Figures S20, S21). We also perform a neighborhood-based `milo` analysis on the immune cells, where we mainly find evidence for increased abundance of the CD8 cell types in the *BRCA1* -carriers as compared to average risk donors (Figure 6c). However, we do not find evidence for the CD4 cell types nor do any cell types have substantial evidence when comparing *BRCA2* -carriers to average risk donors. Several of the `milo` neighborhoods could not be linked to a single cell type, and are assigned the ‘Mixed’ label, which may be a possible explanation. We were also unable to construct shared neighborhoods across both comparisons.

When associating immune cell type composition with age and parous status, as in the original manuscript, all methods (`voomCLR`, `LinDA`, `edgeR` and `milo`) find no significant effects at a 5% FDR level.

## 3 Discussion

We developed `voomCLR`, a statistical method for the differential analysis of cell compositions, and apply it to simulated and real datasets. It is shown that both the CLR transformation and bias correction are crucial aspects in modelling cell compositions. Accounting for the uncertainty involved in the bias correction generally improves false positive control. Additionally, the method’s performance is superior in low sample size settings, where it benefits from accounting for heteroscedasticity and adopting empirical Bayes shrinkage of the residual variances.

While our results show that accounting for compositionality is crucial in the analysis of cell compositions, we also note that non-compositional models originally developed for differential expression analysis, `edgeR`, `DESeq2` and `limma-voom`, perform well in settings with a high sample size and high number of cell types. The latter is expected, as compositional effects get diluted with a high number of features, as is the case in gene expression studies where thousands of features are measured and analyzed simultaneously.

Albeit the good performance, one needs to be careful when applying the methodology when the number of features (cell types) are limited. The CLR transformation can not be expected to work well if the geometric mean is calculated on a very low number of data points, e.g., three cell types. Especially if one or two of these cell types are differentially abundant, it will have a large impact on the geometric mean calculation, which will no longer serve as a reliable reference to compare against. Second, the empirical weights calculation to account for heteroscedasticity relies on a reasonably large number of cell types in order to have a stable lowess fit. This challenge, however, can be tackled via analytical calculation of the weights based on the Delta method. This seems to help especially with false discovery rate control, as shown in our simulations. Lastly, the bias correction assumes most cell types are not differentially abundant. This assumption will be more easily violated with a low number of cell types, although performances are still reasonable when the assumption is violated (Supplementary Figure S6). Even so, the uncertainty in the estimation of the bias term increases as the number of cell types decreases. This also applies for the estimation of the variance of the mode as obtained via the bootstrap.

We have shown the added value of accounting for the uncertainty in the estimation of the bias term, where a procedure is implemented using either a non-parametric or parametric bootstrap procedure. In our evaluations, the non-parametric bootstrap tends to render conservative results, especially in the case of a low number of cell types. The parametric bootstrap has the additional advantage of also accounting for the covariance, and in general performs better than the non-parametric bootstrap. Ultimately, we provide both options in the software, and the decision should be guided by the desire of how strict one may want to control false positive results. Alternatively, when the statistical analysis is aimed at discovery of patterns to be validated in future experiments, the analyst may choose not to use the bootstrap to increase power, and filter out false positives in downstream validation.

We have applied the method on single-cell RNAseq studies, but its application domain should not be limited to this data type. We expect `voomCLR` to be equally useful for differential abundance testing in flow or mass cytometry and microbiome studies.

## 4 Methods

### 4.1 Variance of centered log-ratio transformed counts

As before, let *Y*_*ip*_ denote the population counts for cell population *p* ∈ {1, …, *P* } in sample *i* ∈ {1, …, *n*}. If we assume that **Y**_*i*_|*N*_*i*_, ***π***_*i*_ ∼ *Mult*(*N*_*i*_, ***π***_*i*_), with **Y**_*i*_ the *P* -dimensional vector of population counts, 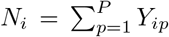 the total number of cells for sample *i* and ***π***_*i*_ the *P* -dimensional vector of expected relative abundances. Then, this can be approximated by assuming independent Poisson distributions for each population, i.e., *Y*_*ip*_ |*λ*_*ip*_ *Poi*(*λ*_*ip*_), where *λ*_*ip*_ = *π*_*ip*_Σ_*p*_*λ*_*ip*_ and 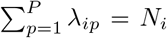. First, note that we can rewrite the CLR transformation as

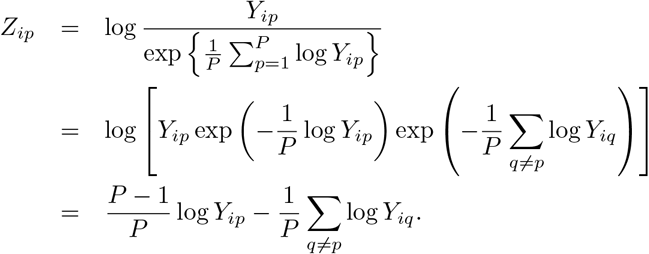

The variance of CLR-transformed counts can be approximated using the Delta method [Dorfman, 1938, Doob, 1935] based on the Poisson assumption.

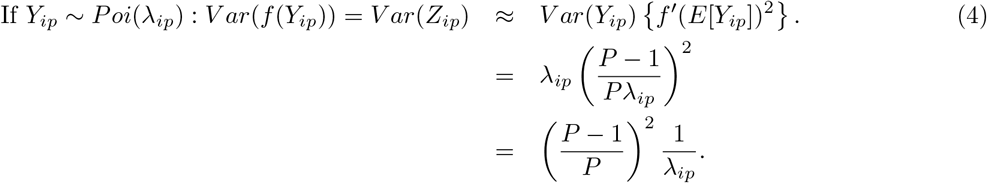

The Multinomial assumption may be too restrictive, therefore several methods rely on a Dirichlet-Multinomial distribution to model cell counts. In turn, the Dirichlet-Multinomial distribution across cell populations can be approximated by assuming a negative binomial distribution for each cell population, i.e., *Y*_*ip*_ ∼ *NB*(*µ*_*ip*_, *ϕ*_*p*_). Based on this, the Delta method would then approximate the variance as

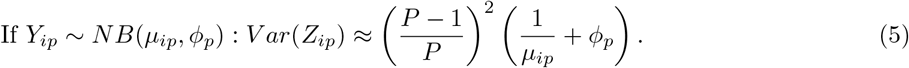

### 4.2 Bias correction when modeling transformed counts

Consider a log-linear model on absolute abundances *X*_*ip*_

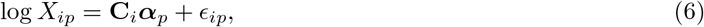

with ***α***_*p*_ the *c* × 1 vector of regression coefficients and **C**_*i*_ being the row corresponding to sample i of the *n* × *c* design matrix **C**, used for representing the experimental covariates taken into account. In a DA analysis, we are interested in testing the null hypothesis that (a linear combination of) ***α***_*p*_, say *α*_*jp*_, is equal to zero, i.e., *H*_0_ : *α*_*jp*_ = 0 for every *p* ∈ {1, …, *P* }.

We usually do not have access to the absolute abundances *X*_*ip*_ but instead must work with the observed cell type counts *Y*_*ip*_. Since these data only contain relative abundance information, one may consider a CLR-transformation, i.e.,

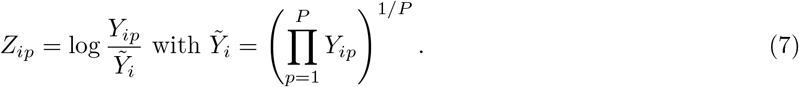

We could similarly fit a linear model on the transformed data,

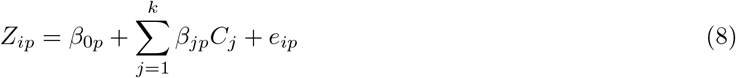

However, regression coefficients ***β*** from this linear model are biased with respect to the coefficients from Equation (6) [Zhou et al., 2022]. Indeed, Zhou et al. [2022] show that 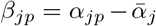,with 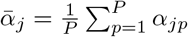. Since *α*_*jp*_ and 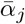 are unidentifiable given an estimate for 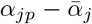, Zhou et al. [2022] make the assumption that most *α*_*jp*_ = 0, or more precisely that the mode of the distribution of *α*_*jp*_ across populations, equals zero. Given this assumption, they estimate 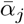 by shifting the distribution of our estimates of 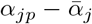 such that it has a mode at zero. The shift is our estimate for 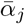,hence providing us with the bias correction term. Concretely, we calculate the bias-corrected coefficient

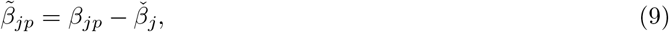

with 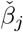 the mode of the *β*_*jp*_ parameters from Model (8), calculated across cell types.

#### 4.2.1 Accounting for the uncertainty of estimating the bias correction term

The variance on 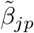 is

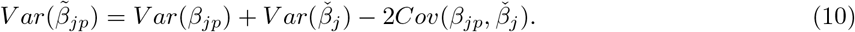

Zhou et al. [2022] argue that as the number of samples and the number of features tend to infinity, *V ar*(*β*_*jp*_) dominates 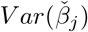 and 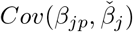 under mild conditions, and therefore rely only on *V ar*(*β*_*jp*_) for statistical inference on 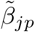.However, in the context of modeling cell type composition this is an unrealistic argument as the number of cell types (features) is usually limited. We therefore attempt to provide a closer approximation to the variance of the bias-corrected parameter by accounting for additional uncertainty involved in estimating the bias correction term,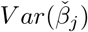. For each (linear combination of) parameter(s) of interest, say **L**^*T*^***β***_*p*_, with **L** a *c* × 1 vector specifying the linear combination of interest, a non-parametric bootstrap procedure [Efron, 1979] is adopted by resampling *β*_*jp*_ across *p*, with replacement, and recalculating **L**^*T*^***β***_*p*_ as needed. By default, *B* = 4000 resamples are taken and, for each *j* ∈ {1, …, *c*}, the dimension of the bootstrapped ***β***_**j**_^*^ is the same as for the original vector ***β***_**j**_, i.e., *P* × 1. For each bootstrap sample and each (linear combination of) coefficient(s), we calculate the mode 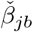 and we approximate the variance of the bias term by

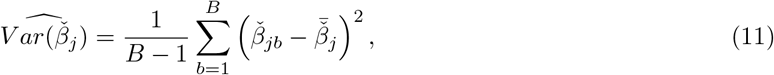

with 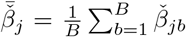. If interest lies in testing the null hypothesis that 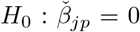, then this would correspond to

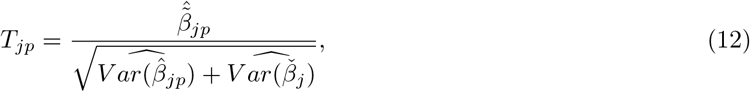

where 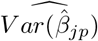 is the estimated moderated variance on 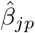 from `limma` [Smyth, 2004].

In Supplementary Methods, we describe an analogous parametric bootstrap procedure, which additionally allows to take into account the covariance term in Equation (10).

### 4.3 voomCLR software implementation

A software implementation of the proposed methodology has been developed as R package. The code required for running a cell composition analysis is similar to a limma-voom analysis, as our implementation reuses functions and code chunks from the `limma` software [Smyth, 2004]. In particular, in `voomCLR`, we adapt the original `voom` function for estimating the mean-variance trend to adopt the CLR transformation and calculating the mean-variance trend for CLR-transformed counts. Model fitting, setting up contrasts and empirical Bayes shrinkage of the residual variance occurs via the native `lmFit`, `contrasts.fit` and `eBayes` functions from `limma`. For statistical inference, we leverage the code from `limma`’s `topTable` function and create a new function, `topTableBC`, which internally implements the bias correction as implemented in the `linDA` software and allows propagating bias correction uncertainty via bootstrapping.

The original `voom` implementation estimates the mean-variance trend empirically, using a lowess trend [Law et al., 2014], which is then leveraged to calculate heteroscedasticity weights to be used in linear model fitting. This is not robust with a low number of features (in our case, cell types). The `voomCLR` function allows for an alternative estimation of heteroscedasticity weights by analytically calculating them using the Delta method, using the derivations shown in Methods. The user has the option to choose between variance approximation based on the Poisson or negative binomial approximation.

The software package is publicly available on GitHub at https://github.com/koenvandenberge/voomCLR and will be submitted to Bioconductor.

### 4.4 Simulation frameworks

#### 4.4.1 Parametric simulation using a Dirichlet-Multinomial distribution

The Multinomial distribution is a natural choice for modeling compositional cell population counts in various contexts. Nevertheless, it falls short in fully explaining the inherent variability observed between biological replicates, a phenomenon known as overdispersion. To address this issue, it is common to treat the Multinomial probabilities as random variables drawn from a Dirichlet distribution to effectively capture the overdispersion. In this study, compositional count samples are generated using a hierarchical Dirichlet-Multinomial distribution. This allows for a flexible and realistic simulation of compositional cell population data, accommodating the inherent variability in biological replicates more accurately. Below, we outline the procedure used for the simulation.

The objective is to simulate compositional cell count data for *P* populations (cell types) measured on two independent groups of samples with replicate size *n*_1_ and *n*_2_. Let **Y**_*i*_ be a *P*-dimensional vector, representing the count vector of a sample *i* ∈ {1, …, *n*}. We assume **Y**_*i*_ ∼ 𝒟ℳ (***π***_*i*_, *θ*_*i*_, *N*_*i*_), where ***π***_*i*_ and ***θ***_*i*_ are *P* - dimensional vectors of Multinomial probabilities of sample *i* and Dirichlet parameters, respectively. For population *p*, we define the Dirichlet parameter 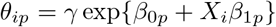, where *X*_*i*_ = 1 if sample *i* is from group 2 and 0 if from group 1. Note that the Dirichlet parameters are the same for all samples from the same group. We denote *v*_0_ and *v*_1_ to be two sets of cell types (|*v*_0_| + |*v*_1_| = *P* and *v*_0_ ∩ *v*_1_ = θ) for which the null and alternative hypothesis holds, respectively. Note that for a population *p* ∈ *v*_0_, its expected absolute count is the same between the two groups of samples. The parameter 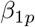 controls the magnitude of differential abundance between the two groups of samples for population *p* ∈ *v*_1_ and the multiplication factor, *γ* > 0 controls the level of variability among samples, which is set constant for both sample groups. A low value of *γ*, results in large variability between samples. The vectors ***β***_**0**_ and ***β***_**1**_ are configured to exhibit variation across cell populations by being sampled from normal distributions. Specifically, 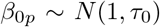, where *τ*_0_ governs the variability between cell populations in terms of abundance levels. In our simulation study, we chose *τ*_0_ = 2 to generate cell population data that mirrors the variability observed in the Lupus case study dataset. Similarly, for 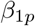, we employed 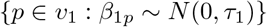, where *τ*_1_ = 2 to simulate effect sizes.

Given ***θ***_*i*_, the Multinomial probabilities for sample *i* are simulated from a Dirichlet distribution. That is, ***π***_*i*_|***θ***_*i*_ ∼ 𝒟 (***θ***_*i*_). Subsequently, for sample *i*, using the obtained ***π***_*i*_ and the desired total number of cells *N*_*i*_, the count vector **Y**_*i*_ is sampled from a Multinomial distribution. That is, **Y**_*i*_|***π***_*i*_, *N*_*i*_ ∼ ℳ (*N*_*i*_, ***π***_*i*_). The simulated data can be denoted by an *n* × *P* matrix **Y**. Data were generated under the following scenarios:

1. Low (*γ* = 1.5), medium (*γ* = 1) or high (*γ* = 0.25) levels of variability between samples by controlling the, parameter.
2. Different sample sizes *n*_1_ = *n*_2_ ∈ {5, 10, 20}.
3. Different number of cell types *P* ∈ {5, 10, 20, 50, 100}.
4. Different number of truly differentially abundant cell types (pDA): 10, 20, 50, 75 and 85% of the number of cell types (*P*)

In most of the simulation settings, the number of truly differentially abundant populations | *v*_1_| is set to be 20% of *P* (rounded to the nearest higher integer number) unless specified otherwise.

#### 4.4.2 Non-parametric simulation framework

Although simulation from Dirichlet-Multinomial distribution provides flexibility to simulate a wide range of scenarios, the parametric assumptions inherent in modeling cell count data may not always align with reality. In recognition of this limitation, and to steer clear of such parametric assumptions, we adopt a novel distribution-free approach. This alternative method begins with real cell population data, enabling the simulation of realistic new datasets with a built-in truth. Below, we outline the procedure.

This simulation method uses real data as input. We start with a set of samples between which we expect no systematic biological signal, for example, a group of healthy patients from the lupus cell population dataset. In brief, the simulation method works by first randomly splitting the samples (patients) into two equally sized groups. Again, we expect no systematic signal between these groups. The introduction of signal happens by replacing the cell counts of a randomly selected population with the cell counts of another randomly selected population, within each sample for one of the two artificially created groups. In order to simulate the compositional effect without changing the total number of cells for each sample, we make a compositional correction for all other populations that were not changed.

More formally, let **Y** be the *n* × *P* cell population count matrix that serves as the basis for the simulation, with elements *Y*_*ip*_, *i* ∈ {1, …, *n*}, *p* ∈ {1, …, *P* }. We randomly split the *n* samples into two mutually exclusive groups of size *n*_1_ and *n*_2_, where *n*_1_ + *n*_2_ ≤ *n*. Let 𝒢_1_ and 𝒢_2_ denote the set of samples belonging to group 1 and group 2, respectively. Let 𝒱= {(*p, q*) : *p, q* ∈ {1, … *P*}, and *p* ≠ *q* be a set of indices for a pair of cell populations.

1. Randomly select 𝒱. Let **y**_*p*_ and **y**_*q*_ denote the *n*_2_-dimensional vectors of cell counts for all samples in 𝒢_2_ for populations *p* and *q*, respectively. Additional notes on the selection of 𝒱 are given below.
2. Cell count replacement/swap. From the set 𝒱, population *p* was selected to have signal introduced, and the counts are replaced with those of population *q* in the second group, i.e., **y’**_*p*_ = **y**_*q*_, where **y’**_*p*_ is the new cell count vector of population *p*.
3. Calculate spillover cell count: Let **d**_*p*_ = **y’**_*p*_ **y**_*p*_ is an *n*_2_-dimensional vector denoting the difference in the cell counts of population *p* introduced by the count replacement in Step 2.
4. Calculate compositional effect: For all other populations *k* ≠ *p*, we calculate the compositional effect caused by replacing population *p* by population *q* as ***c***_*k*(*p*)_ =− **w**_*k*(*p*)_ ⊙**d**_*p*_, where **w**_*k*(*p*)_ is the wight vector with element *w*_*k*(*p*)*i*_ for sample *i* ∈ 𝒢_2_ defined as 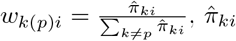 is the estimated fraction of cell population *k* in sample *i*, and ⊙ denotes element-wise multiplication. See the notes below for additional details.
5. Correct for compositional effect. Add the compositional effect for every cell population *k* ≠ *p*, i.e., 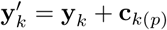.
6. Repeat steps 1-5 until a desired number of cell populations is set to be differentially abundant.

*Note 1:* In Step 1, two populations can be selected randomly but to simulate realistic data, we select pairs of populations with probability sampling. The sampling probability for a pair of cell populations p and q is inversely proportional to the Euclidean distance between **x**_*p*_ and **x**_*q*_, where **x** is the CLR transformed vector.

*Note 2:* In Step 4, the weights are introduced to redistribute excess counts to other cell populations proportional to their relative abundance. This, for example, keeps a rare population being rare in the simulated data.

One may consider swapping, for instance, two cell populations within the second group of samples to introduce a signal. However, the objective is to simulate realistic cell population data in a manner where the introduction of differential abundance signals in a specific cell population systematically influences the abundance of other cell populations. This process should ultimately emulate the natural data-generating mechanism of cell population data.

The non-parametric simulation is conducted using one of the two real datasets as baselines: (1) all biological replicates from the healthy patients in the lupus dataset from processing cohort 4, or (2) all biological replicates from the healthy groups in the Breast Cell Atlas data. Unless specified, a differential abundance signal is introduced to 20% of the cell populations in each simulated dataset.

#### 4.4.3 Performance assessment metrics

The simulated data contain built-in truth about the truly differentially abundant cell populations. This helps to quantify the actual false discovery rate (FDR) and true positive rate (TPR) for a given nominal level of FDR. The actual FDR is defined as the average false positive fraction among the positives (FP/(FP+TP)) across the simulation runs. The TPR is defined as the average true positive proportions (TP/TP+FN)), where FN, FP, and TP denote, respectively, the numbers of false negatives, false positives, and true positives. FDR-TPR curvesare calculated using the iCOBRA Bioconductor R package [Soneson and Robinson, 2016].

### 4.5 Lupus scRNA-seq data case study

The lupus scRNA-seq dataset [Perez et al., 2022] was downloaded from the Gene Expression Omnibus using accession number GSE174188. We use the annotation from the paper to create the cell population count matrix, consisting of 11 cell populations and 355 samples. We remove four samples from African American patients and 3 samples from Hispanic patients, focusing the analysis on the remaining samples from Asian (n=137) and European (n=211) ancestry. Out of the remaining 348 samples from 256 individuals, 188 individuals were sequenced once, while 49 individuals were sequenced twice, 14 individuals were sequenced three times, and 5 individuals were sequenced four times. The sequencing happened in multiple phases, denoted as four different processing cohorts in the original manuscript. We perform a principal components analysis (PCA) on the cell count matrix based on Aitchison’s distance (i.e., Euclidean distance on CLR-transformed counts), respecting the compositional nature of the counts. When modeling the counts, we use the same design matrix for all methods, which includes a fixed effect for processing cohort and main and interaction effects for disease status and ancestry. We account for repeated measurements through repeated sequencing by using a random subject effect in the negative binomial model and `LinDA`. For `voomCLR`, we approximate the same random effect using the `duplicateCorrelation` functionality of `limma` [Smyth, 2004]. Statistical inference is focused on the fixed effect part of the model, where we test for disease effect for European ancestry, disease effect for Asian ancestry, and the interaction effect.

### 4.6 Breast cell atlas case study

The breast atlas case study dataset was downloaded from the cellxgene portal link that was provided in the manuscript [Reed et al., 2024]. Cells belonging to clusters ‘Doublet’, ‘Stripped nuclei’, ‘DDC1’ and ‘DDC2’ (donor-derived clusters 1 and 2) were removed. The immune cells were obtained by first selecting the ‘Stroma enriched’ samples and then subsetting to the immune cell types. As in the original manuscript, the associations of age and parity were done on the mammoplasty samples. These tests were always performed with both variables in the model (e.g., the analysis testing for age also included parity as a variable in the model). Samples with no information on parity status were removed. When comparing high-risk with average risk donors, we also condition on age and parity. Analysis was performed separately on the samples consisting of {AR, HR-BR1} and {AR, HR-BR2} data subsets. In the `milo` analysis of the high risk versus average risk comparison, we were unable to use shared neighborhoods across these comparisons for the downstream statistical inference, since the software errored in the inference evaluation.

### 4.7 Other methods

#### 4.7.1 Negative Binomial GLM (NB_GLM)

Cell abundance counts are used as a response in a negative binomial generalized linear model (NB_GLM), using the `glmmTMB` package (v1.1.9) [Brooks et al., 2017]. An offset corresponding to the logarithm of each sample’s total number of cells is used.

#### 4.7.2 edgeR

Similar to `NB_GLM`, `edgeR` (v3.38.4) is used to fit negative binomial models to the cell abundance counts of each cell type [Robinson et al., 2010]. The offset now corresponds to the logarithm of normalized total cell counts, with normalization factors calculated based on TMM normalization [Robinson and Oshlack, 2010]. Dispersion parameters are estimated using the `estimateDisp` function and model fitting happens through `glmFit`. In the simulation study, the Quasi-Likelihood (QL) method (`glmQLFit`) is used for testing differential abundance and dispersion estimates obtained using the `estimateQLDisp` function.

#### 4.7.3 DESeq2

We also use `DESeq2` (v1.36.0) for negative binomial model fitting, where the total cell counts are now normalized according to the `DESeq2` median-of-ratios normalization [Love et al., 2014]. Further, we run `DESeq2` using default settings.

#### 4.7.4 limma-voom (limma-voom)

In `limma-voom`, cell abundance counts are transformed to log_2_ counts-per-million, which are then used as response in a weighted linear model. The weights are calculated based on an empirical mean-variance trend as described in [Law et al., 2014], and residual variances are shrunken using empirical Bayes [Smyth, 2004]. The `limma` package (v3.52.3) is used for the implementation of `limma-voom`

#### 4.7.5 CLR-based linear model (LM_CLR)

In the `LM_CLR` method, we fit linear models for each cell population, using the CLR-transformed cell counts as response variable. Statistical inference is carried out using t-tests.

#### 4.7.6 LinDA

`LinDA` is designed for analyzing microbiome compositional data. It fits linear regression models on centered log-ratio transformed data, identifies bias terms, and corrects these biases using the mode of the regression coefficients. The package `LinDA` (v0.2.0) is used with the default settings.

#### 4.7.7 propeller

`propeller` [Phipson et al., 2022] is a method proposed for testing differential abundance in compositional cell population data. It applies a transformation (using logit or arcsin square root) on cell counts and fits linear models. Empirical Bayes is used for moderated cell type-specific variance estimation. We try both transformations in the simulation study, with all other parameters kept at the default settings. The `fitPropeller` function from the `speckle` (v1.4.0) R package is used.

#### 4.7.8 DCATS

`DCATS` [Lin et al., 2023] is an R software package developed for differential composition analysis in single-cell RNA sequencing (scRNA-seq) data, with a focus on addressing the uncertainty in cell type assignment. It uses a beta-binomial regression model to analyze raw cell counts, considering dispersion between samples. DCATS corrects for misclassification bias using a similarity matrix between cell types. The package `DCATS` (v1.2.0) is applied with its default settings.

### 4.8 Code availability

Code to reproduce the simulations and case studies is available from GitHub at https://github.com/koenvandenberge/voomCLRPaper.

## 5 Acknowledgements

The authors thank Lucas Beerland and Lieven Clement for their advice on software implementation, and Ewoud De Troyer for feedback on initial methodological developments and software implementation.

## Notes

### Competing Interest Statement

The authors are employees of Janssen Pharmaceutica NV.

