## Supplementary Material for "Assessing differential cell composition in single-cell studies using voomCLR"

### 6 Supplementary Methods

#### 6.1 Parametric bootstrap procedure

A linear model on CLR-transformed cell type counts is fitted,

$$Z_{ip} = \beta_{0p} + \sum_{j=1}^{k} \beta_{jp} C_j + e_{ip}, \tag{13}$$

and we are interested in statistical inference on (a linear combination of) the mean parameter(s) $\beta_{jp}$. Let $\beta_{cp} = \mathbf{L}\boldsymbol{\beta}_p$ denote the contrast of interest, where $\mathbf{L}$ is a $1 \times P$ contrast vector and $\boldsymbol{\beta}_p$ the $P \times 1$ vector of regression coefficients. Analogous to the non-parametric bootstrap procedure, we develop a parametric bootstrap to accommodate the uncertainty involved in estimation of the bias correction term $\check{\beta}_c$ for the relevant contrast of interest. For each population $p$, we resample $\hat{\boldsymbol{\beta}}_p$ (by default 4000 times) from a multivariate normal (MVN) distribution

$$\hat{\boldsymbol{\beta}}_p^B \sim MVN(\hat{\boldsymbol{\beta}}_p, \hat{\Sigma}_{\hat{\beta}_p}), \tag{14}$$

where $\hat{\Sigma}_{\hat{\beta}_p} = \tilde{\sigma}^2(\mathbf{C}^T\mathbf{WC})^{-1}$, with $\tilde{\sigma}^2$ the moderated residual variance estimate [Smyth, 2004, Law et al., 2014], $\mathbf{C}$ the $n \times p$ design matrix from the model in Equation (13) and $\mathbf{W}$ a $n \times n$ diagonal weight matrix corresponding to the `voom` heteroscedasticity weights [Law et al., 2014]. For each contrast of interest, we then calculate the corresponding mode in each bootstrap sample, i.e., $\check{\beta}_c$. Estimating the variance across bootstrap samples and statistical inference then proceeds as in the non-parametric bootstrap procedure.

### 7 Supplementary Figures

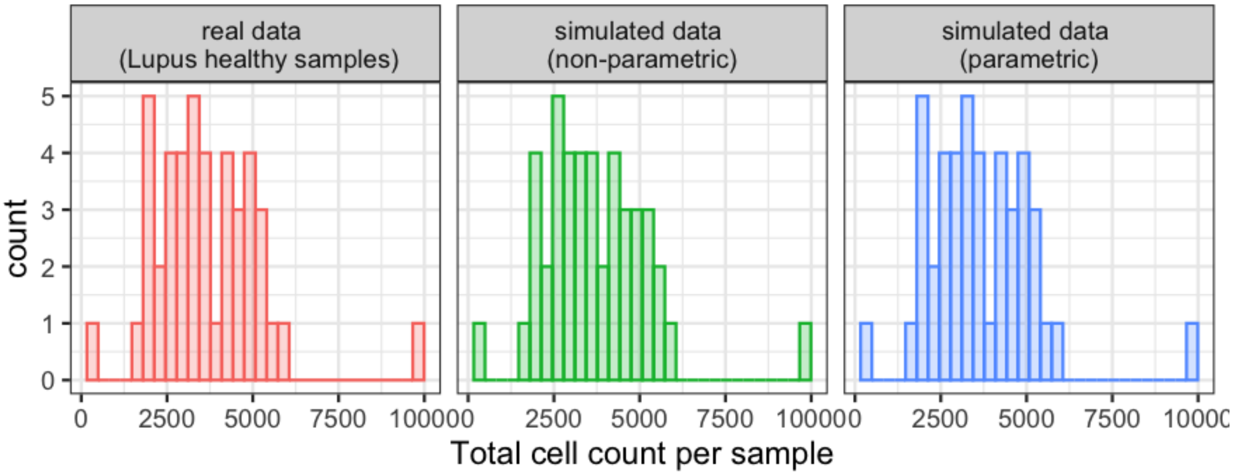


Supplementary Figure S1: *Simulation study:* Comparison of the distribution of the total cell counts per sample between real data (lupus healthy samples from processing cohort 4) and simulated data (**non-parametrically** and **parametrically** simulated data). Note that the distributions from the real and parametrically simulated data are identical because the parametric simulation procedure uses the total cell count from the real data to simulate cell counts from the Dirichlet-Multinomial mixture distribution.

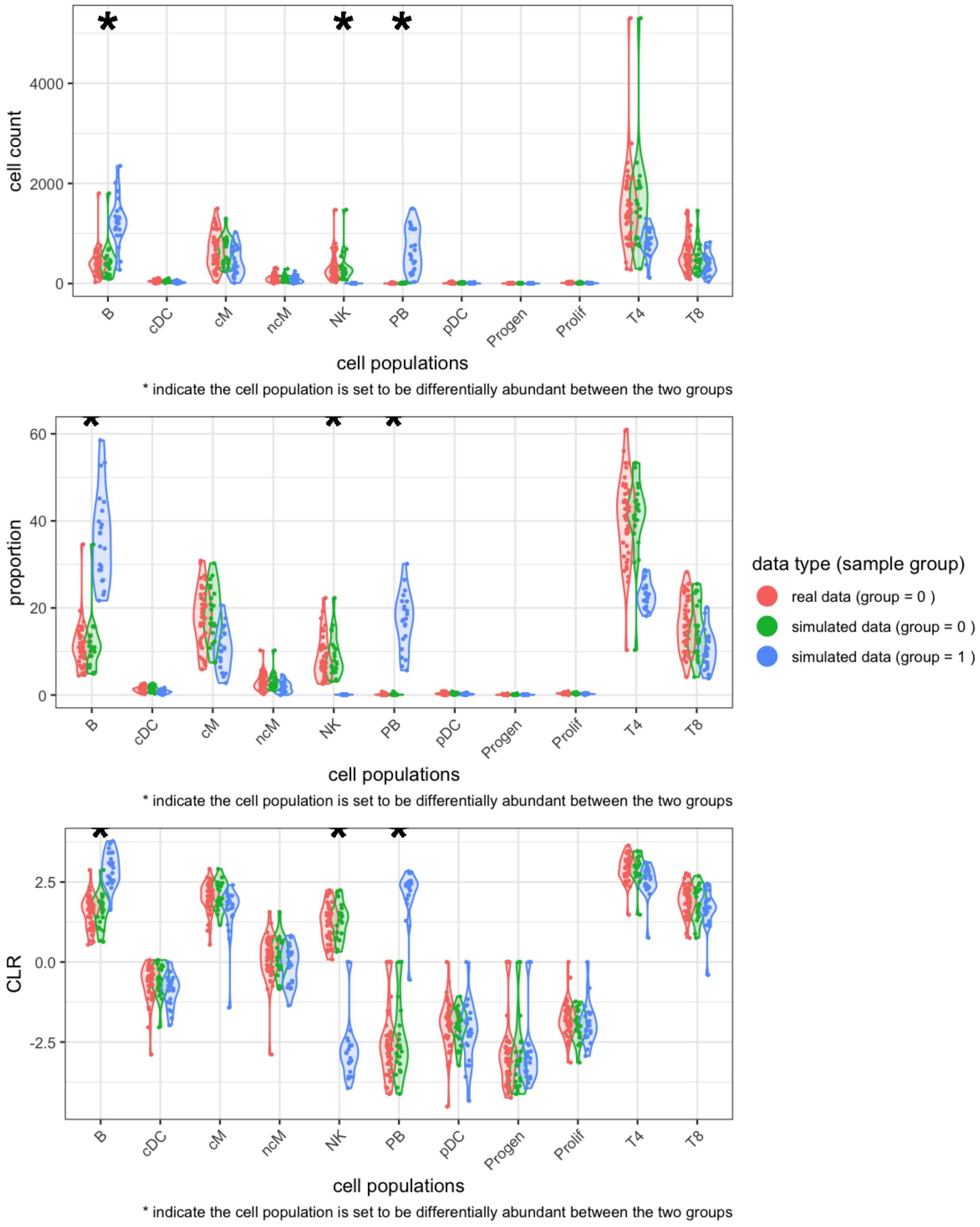


Supplementary Figure S2: *Simulation study:* Comparison of the distribution of the cell counts, proportions, and CLR-transformed counts between real data (lupus healthy samples from processing cohort 4) and **non-parametrically** simulated data.

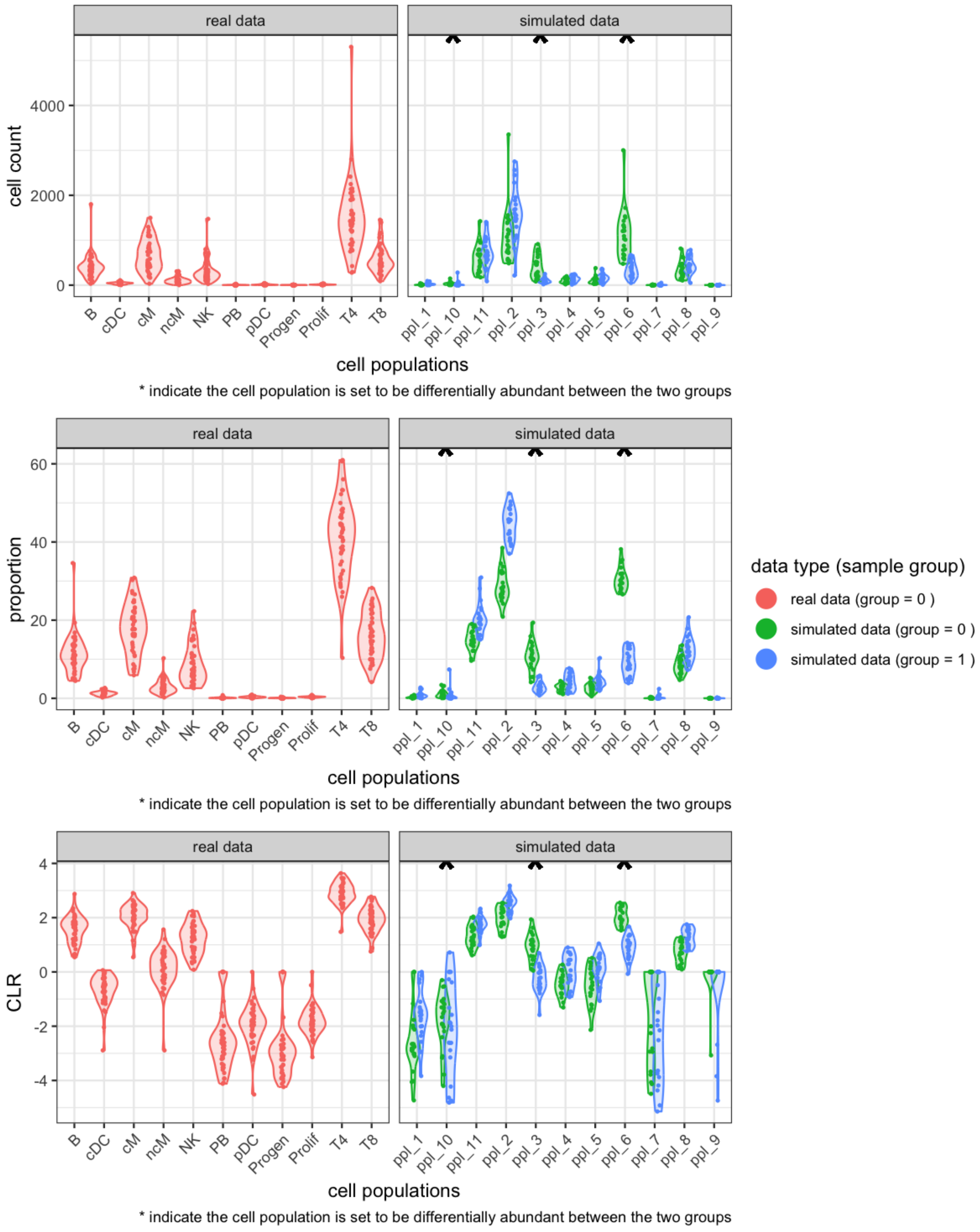


Supplementary Figure S3: *Simulation study:* Comparison of the distribution of the cell counts, proportions, and CLR-transformed counts between real data (lupus healthy samples from processiong cohort 4) and **parametrically** simulated data.

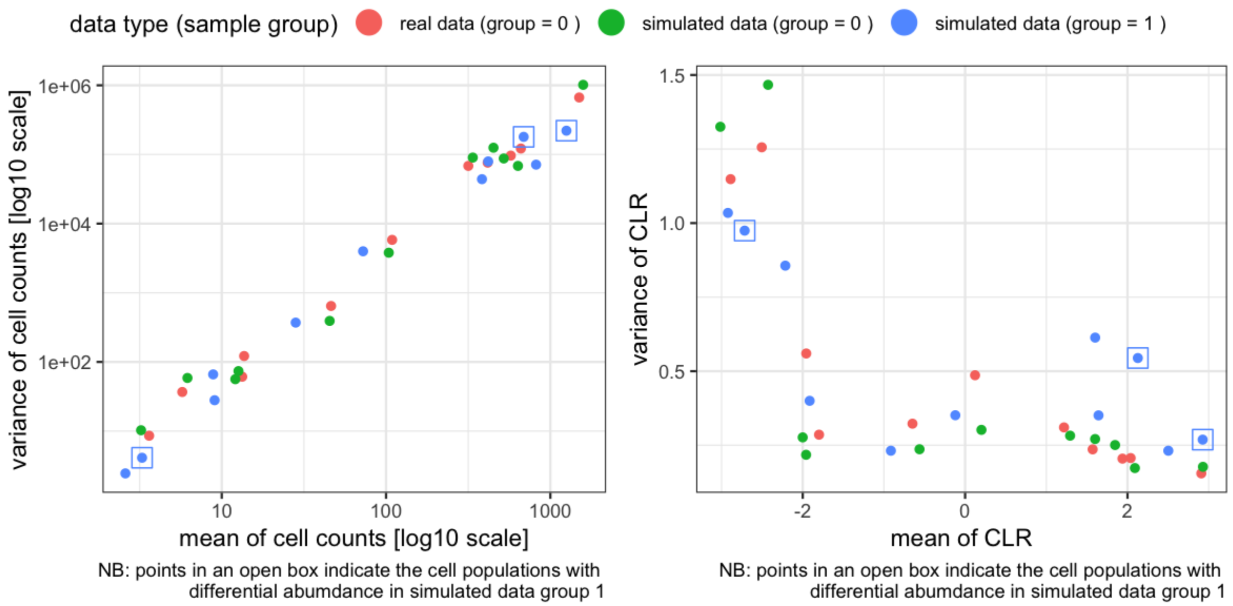


Supplementary Figure S4: *Simulation study:* Comparison of the mean-variance trend for the cell counts and CLR-transformed counts between real data (lupus healthy samples from processing cohort 4) and **nonparametrically** simulated data.

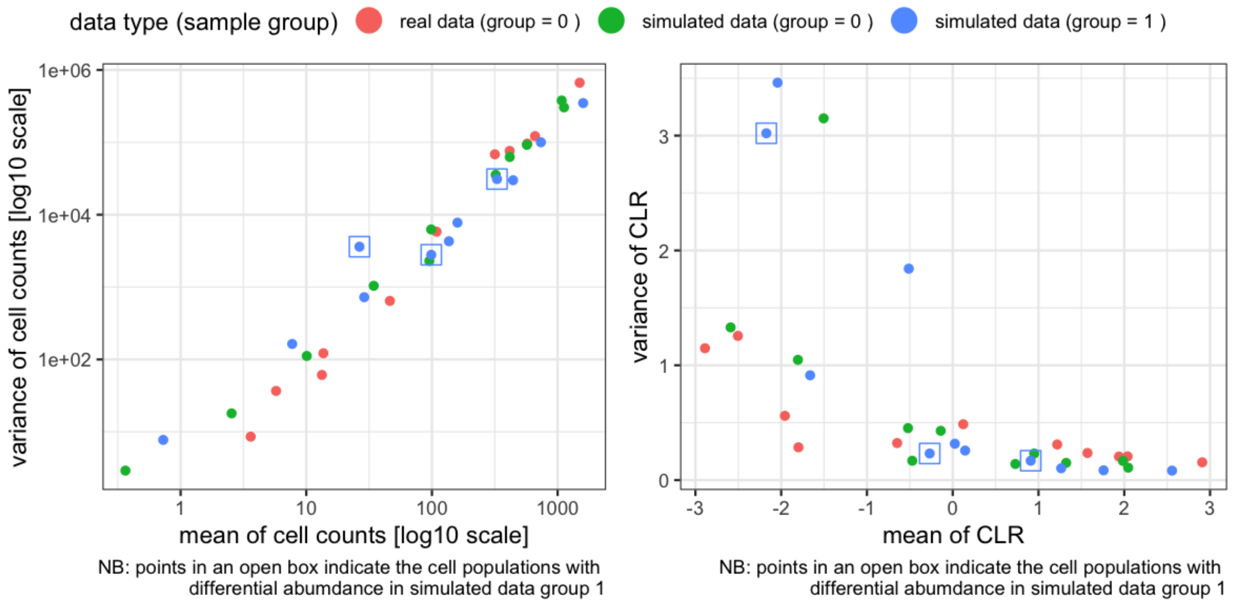


Supplementary Figure S5: *Simulation study:* Comparison of the mean-variance trend for the cell counts and CLR-transformed counts between real data (lupus healthy samples from processing cohort 4) and **parametrically** simulated data.

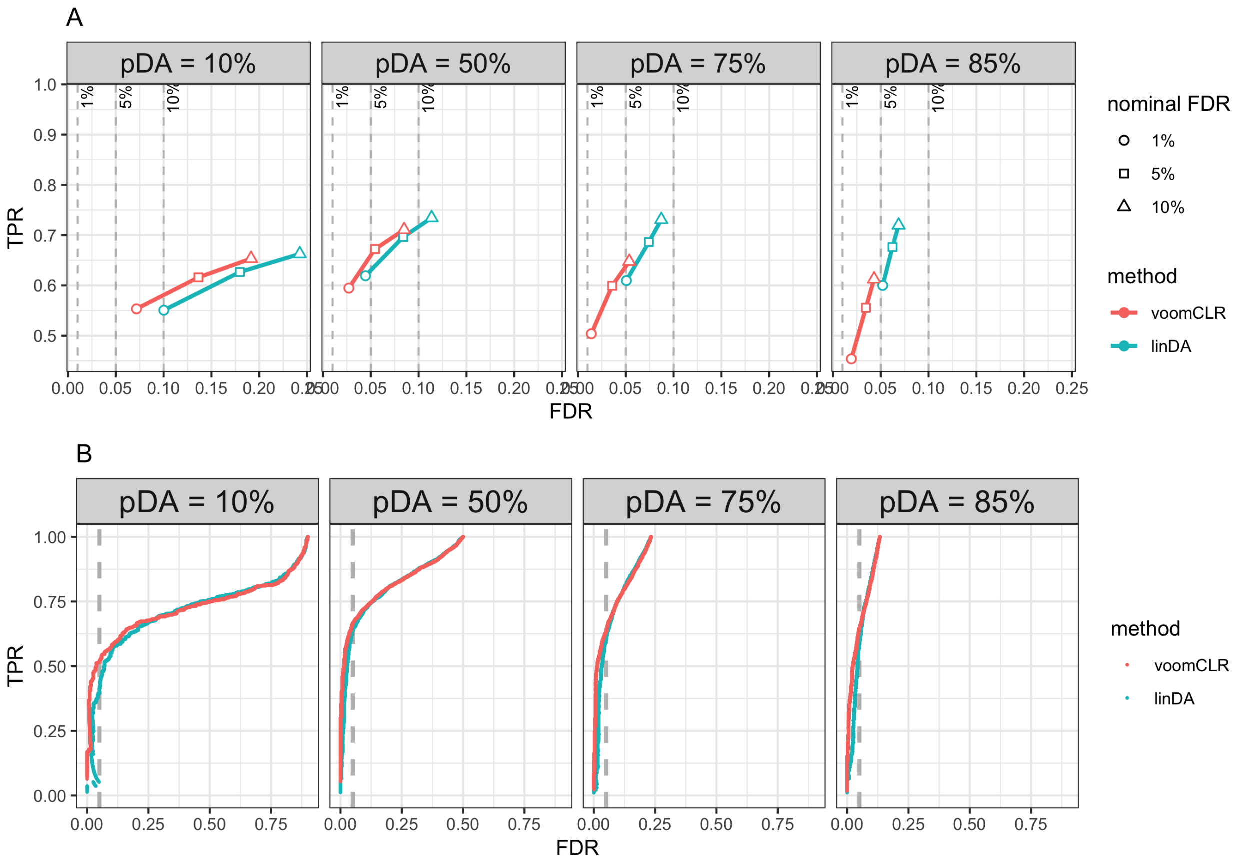


Supplementary Figure S6: *Simulation study:* Performance curves for evaluating the performance of voomCLR and LinDA for in simulation settings with 10, 50, 75 and 85% of the cell populations are set to be DA. The data is simulated from a Dirichlet-Multinomial distribution, with a medium level of variability (Dirichlet parameters scaling factor $\gamma = 1$), and 10 samples per group ($n = 10$). Reported metrics are averages from 250 simulation runs for each scenario. (A) FDR-TPR curves, where the metrics are calculated at 1%, 5%, and 10% nominal FDR, and (B) ROC curves, where the metrics at 5% nominal FDR are shown indicated with the circle.

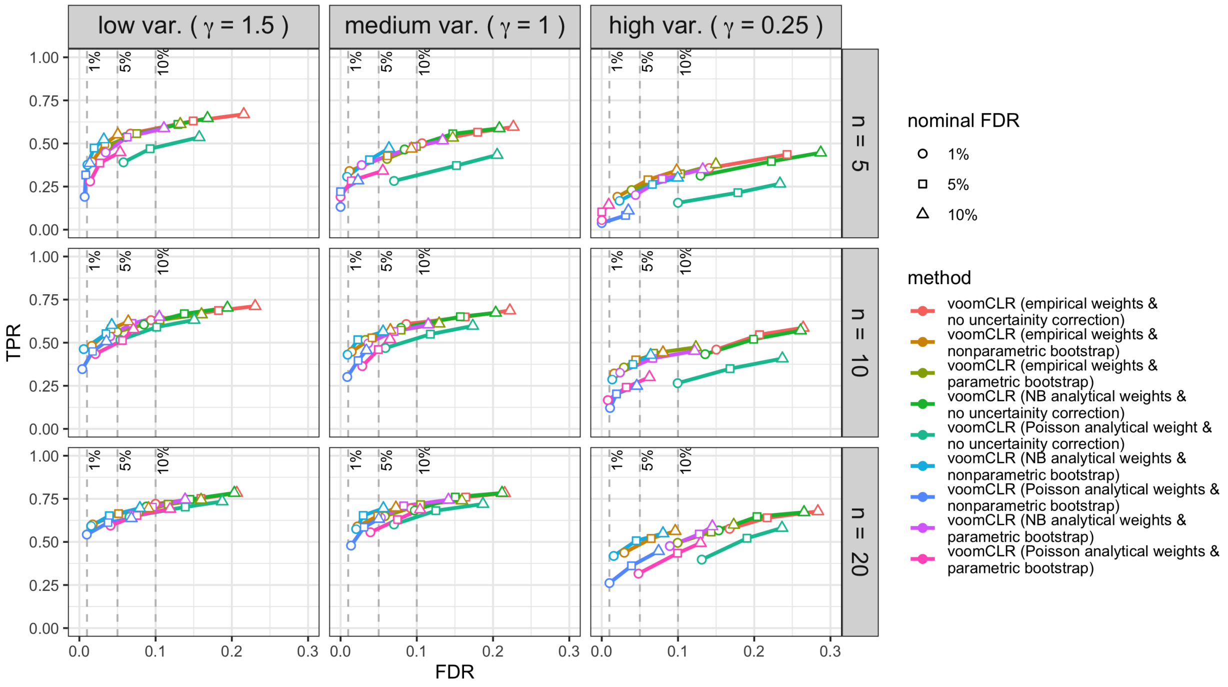


Supplementary Figure S7: *Simulation study:* FDR-TPR curves for evaluating the performance of voomCLR with different configurations in simulated cell population data with three different levels of variability among samples and three different sample sizes. The data is simulated from a Dirichlet-Multinomial distribution, with 3 different levels of variability (low, medium and high) controlled by a scaling factor $\gamma$ for the Dirichlet parameters and 3 different sample sizes per group ($n = 5, 10$ and 20). Reported metrics are averages from 250 simulation runs for each simulation scenario and performance metrics are calculated at $1, 5,$ and 10% nominal FDR. Each simulated data consists of two independent groups of samples for 11 populations.

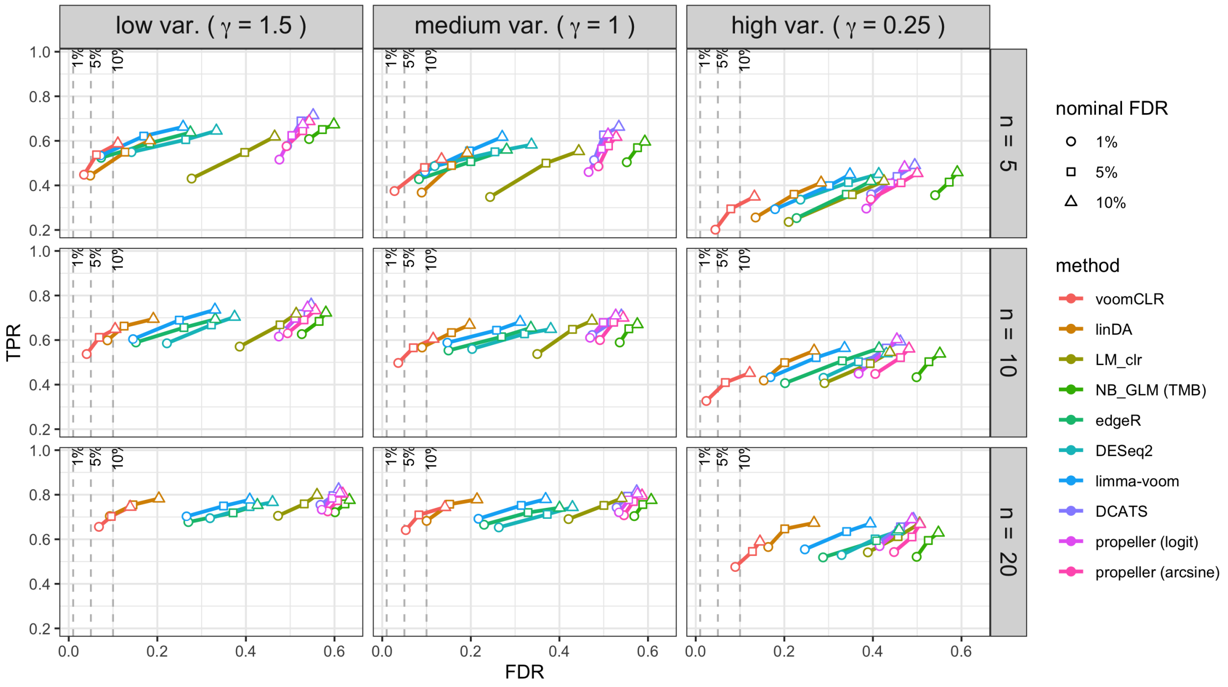


Supplementary Figure S8: *Simulation study:* FDR-TPR curves for evaluating the performance of voomCLR and other methods in simulated cell population data with three different levels of variability among samples and three different sample sizes. The data is simulated from a Dirichlet-Multinomial distribution, with 3 different levels of variability (low, medium and high) controlled by a scaling factor $\gamma$ for the Dirichlet parameters and 3 different sample sizes per group ($n = 5, 10$ and $20$). Reported metrics are averages from 250 simulation runs for each simulation scenario and performance metrics are calculated at $1, 5,$ and $10\%$ nominal FDR. Each simulated dataset consists of two independent groups of samples for 11 populations.

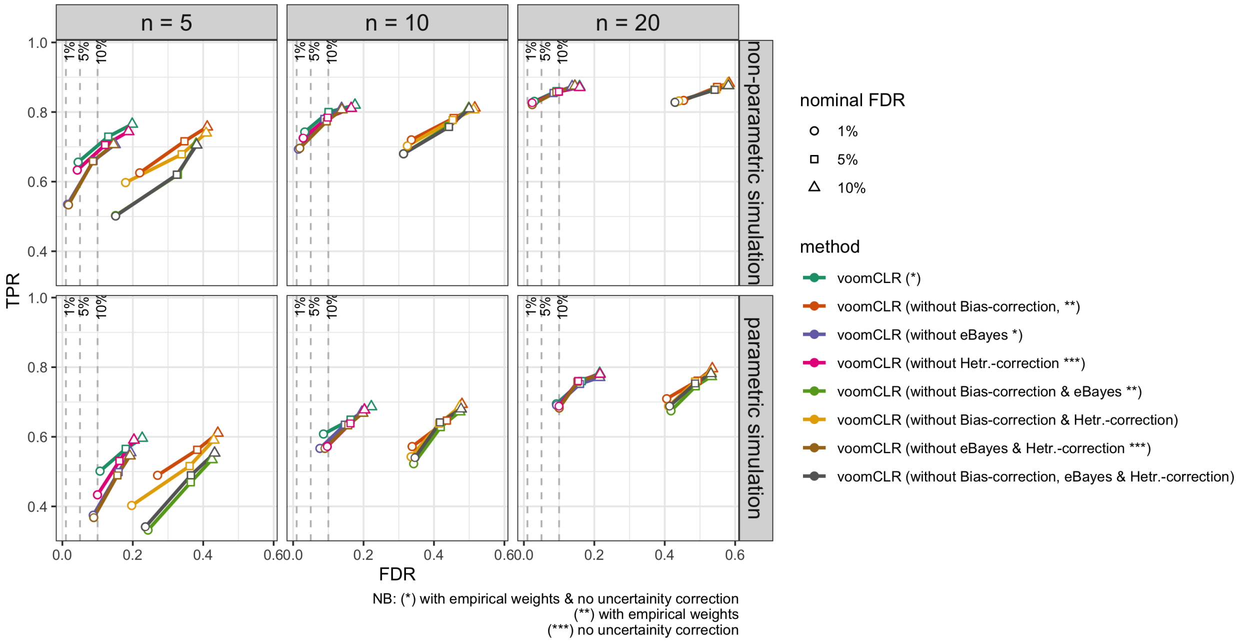


Supplementary Figure S9: *Simulation study:* FDR-TPR curves for evaluating the performance of voomCLR with and without either bias correction, heteroscedasticity or empirical Bayes shrinkage in simulated cell populations with $P = 11$ cell populations. The non-parametric simulation uses the healthy samples from the Lupus dataset in processing cohort 4 as a baseline. The data is simulated using the non-parametric and parametric simulation procedures. Reported metrics are averages from 250 simulation runs for each simulation scenario. The reported performance metrics are at 1%, 5%, and 10% nominal FDR. Notes: annotation in legend "without Bias-correction" is for voomCLR without bias correction, "without eBayes" is for voomCLR without empirical Bayes shrinkage and "without Hetr.-correction" is for voomCLR without accounting for mean-variance trend and all combinations thereoff. Also note that the voomCLR configuration used in these settings is (if applicable) with empirical observational weights and parametric bootstrap procedure.

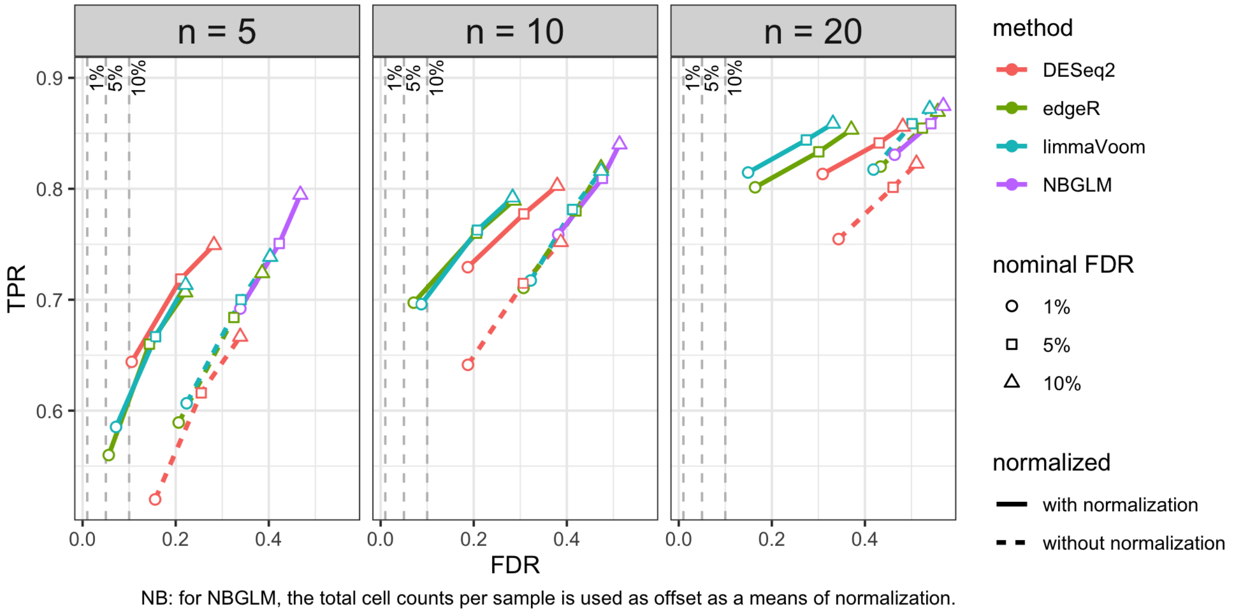


Supplementary Figure S10: *Simulation study:* FDR-TPR curves for evaluating the effect of library size normalization on the performance of edgeR, DESeq2, and limma-voom methods in simulated cell populations with $P = 11$ number of cell populations and 3 different sample sizes per group ($n = 5, 10$ and 20). The data is simulated using the non-parametric simulation procedure using the healthy samples from the Lupus dataset in processing cohort 4 as a baseline. Reported metrics are averages from 250 simulation runs for each simulation scenario. The reported performance metrics are at 1%, 5%, and 10% nominal FDR.

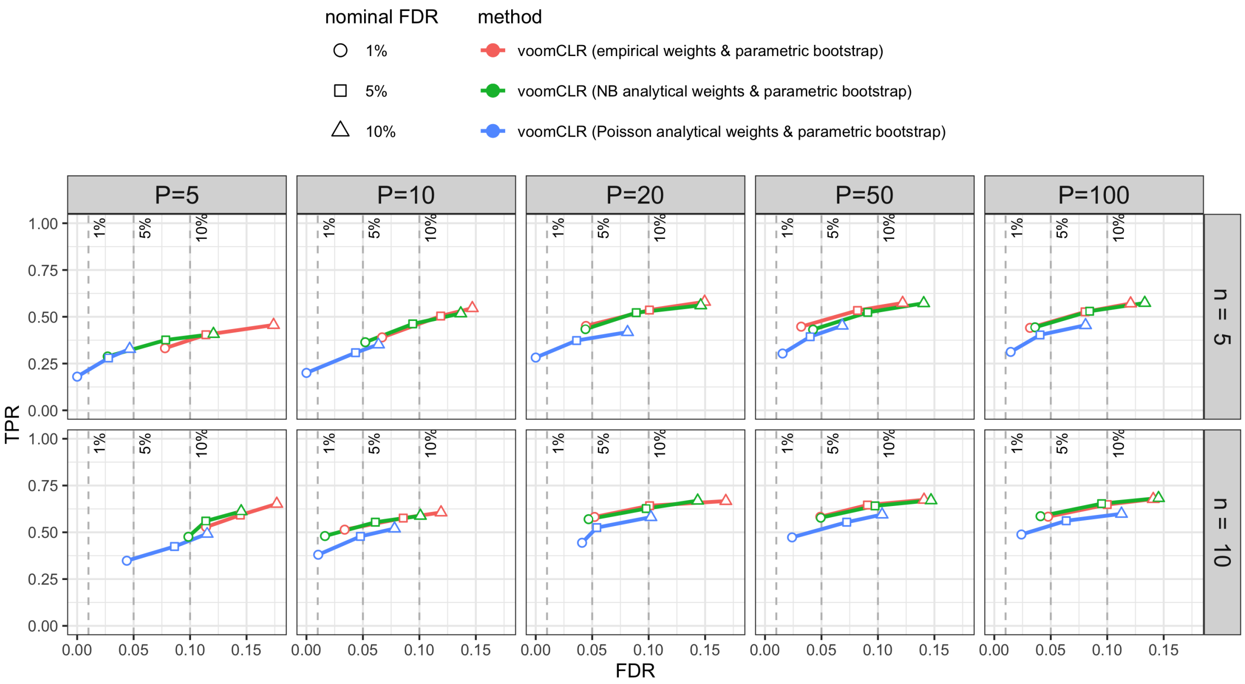


Supplementary Figure S11: *Simulation study:* FDR-TPR curves for evaluating the performance of 3 configurations of voomCLR with respect to estimation of observational heteroscedasticity weights (empirical or analytical weights from negative binomial or Poisson distribution all with parametric bootstrap) in simulated cell population data for 5 different numbers of cell populations ($P = 5, 10, 20, 50$ and 100). The data is simulated from a Dirichlet-Multinomial distribution, with medium level of variability (Dirichlet parameters scaling factor $\gamma = 1$), and 2 different sample sizes per group ($n = 5$ and 10). Reported metrics are averages from 250 simulation runs for each simulation scenario and performance metrics are calculated at 1%, 5%, and 10% nominal FDR.

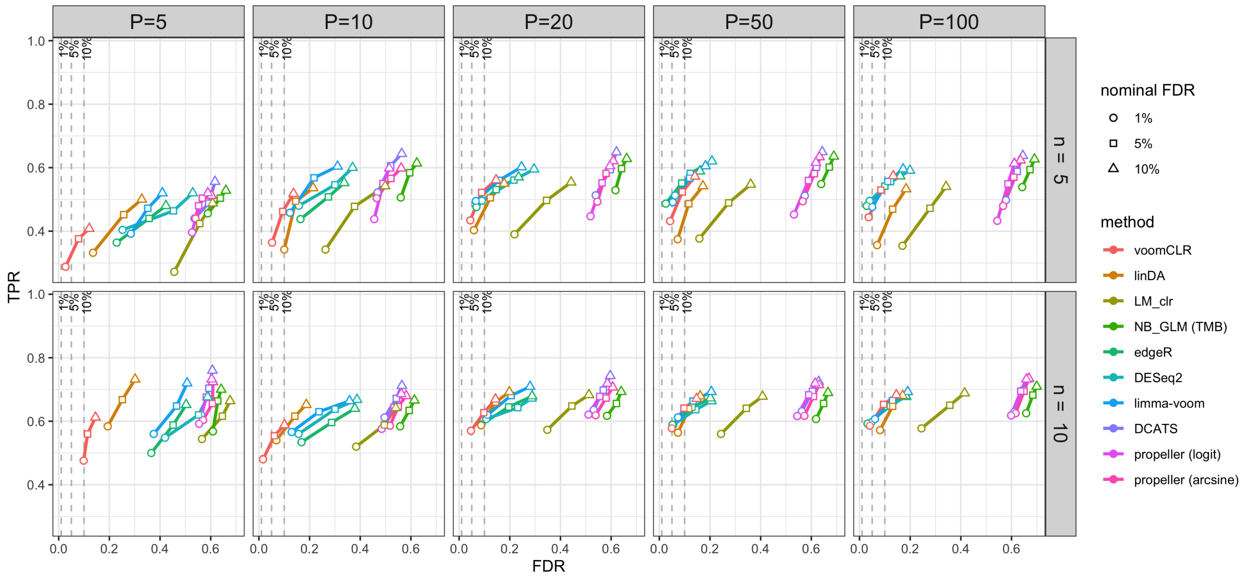


Supplementary Figure S12: *Simulation study:* FDR-TPR curves for evaluating the performance of voomCLR and other methods for 5 different numbers of cell populations ($P = 5, 10, 20, 50$ and 100). The data is simulated from a Dirichlet-Multinomial distribution, with medium level of variability (Dirichlet parameters scaling factor $\gamma = 1$), and 2 different sample sizes per group ($n = 5$ and 10). Reported metrics are averages from 250 simulation runs for each simulation scenario and performance metrics are calculated at 1%, 5%, and 10% nominal FDR.

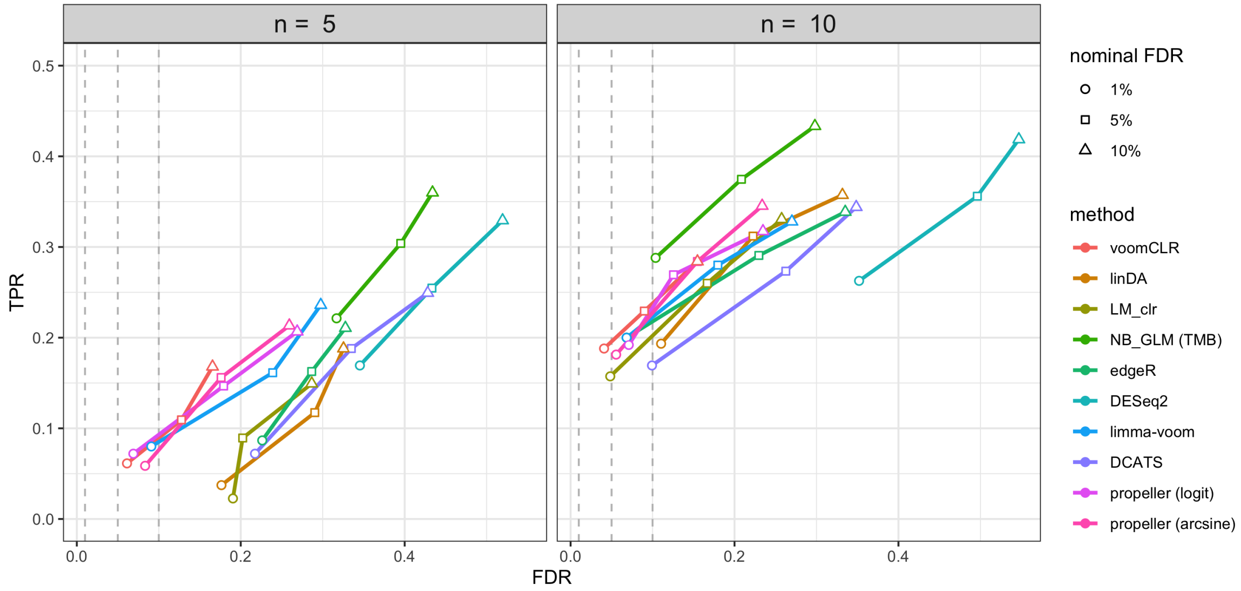


Supplementary Figure S13: *Simulation study:* FDR-TPR curves for evaluating the performance of voomCLR and other methods with non-parametrically simulated cell population count data using the Breast cell atlas data as input (healthy samples are used as the baseline for the simulation). The number of cell populations in the simulated data is 15. Reported metrics are averages from 250 simulation runs for each simulation scenario and performance metrics are calculated at 1%, 5%, and 10% nominal FDR

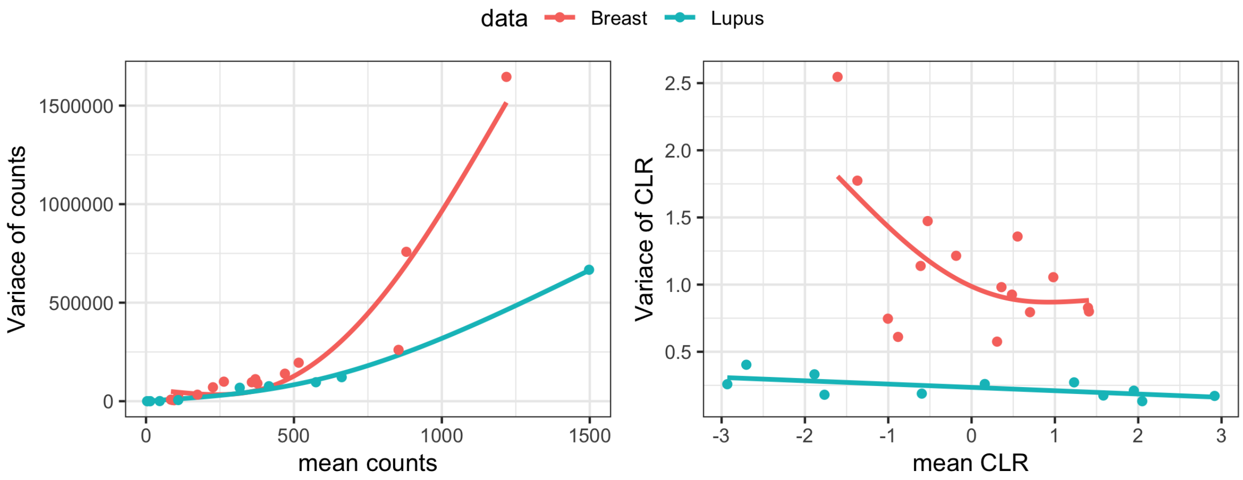


Supplementary Figure S14: *Breast atlas and Lupus case study: comparison of variability.* Mean-variance trend in the Breast atlas (healthy samples) and Lupus (healthy samples in processing cohort 4) datasets. Each point is a cell type and the solid lines are the smoothed trends for the mean variance association of counts (left) and CLR transformed counts (right).

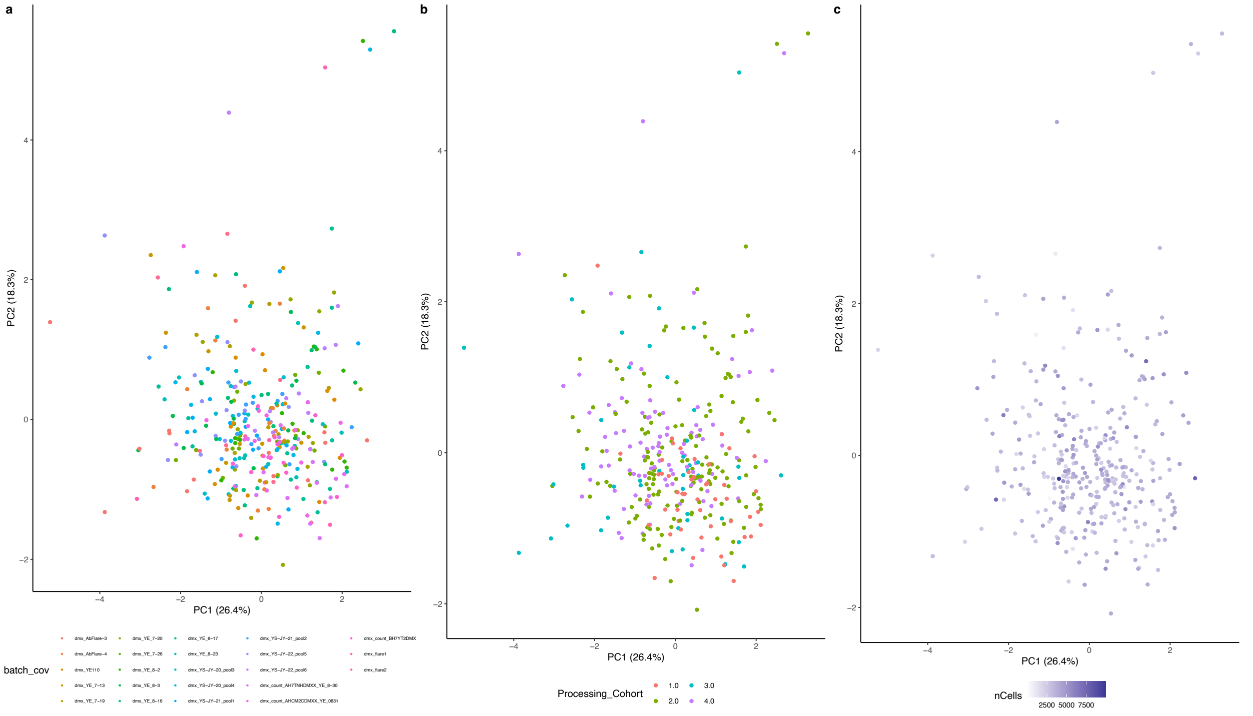


Supplementary Figure S15: *Lupus case study: Scatterplot of first two principal components of Aitchison's distance PCA, coloring by technical variables.* **(a)** Samples colored by batch. **(b)** Samples colored by processing cohort. **(c)** Samples colored by the total number of cells in each sample.

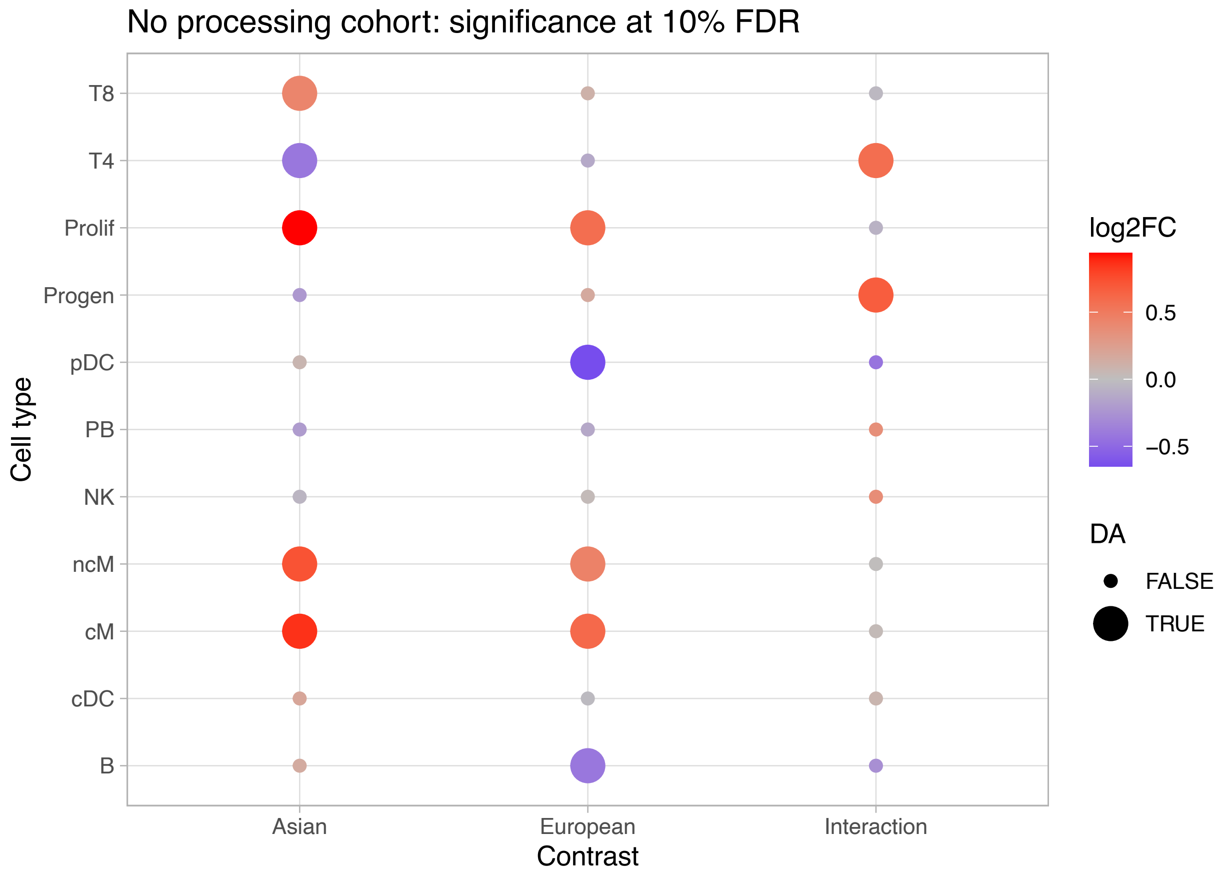


Supplementary Figure S16: *Lupus case study: Heatmap showing* voomCLR *results when not accounting for processing cohort.* Contrasts are denoted in columns and cell types in rows. Each point is colored according to the log2 fold-change, and point size denotes whether the FDR-adjusted p-value is below the 10% level.

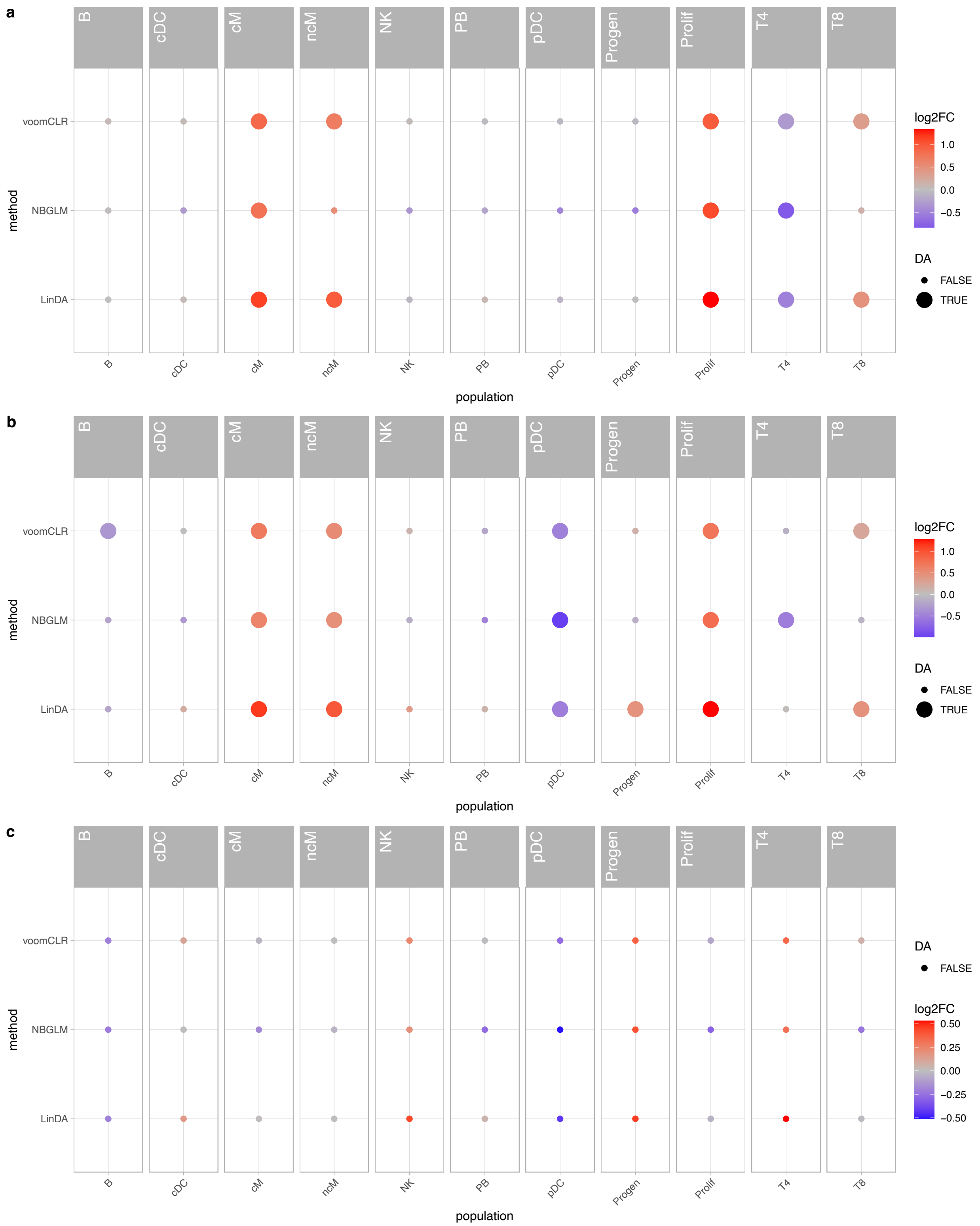


Supplementary Figure S17: *Lupus case study: Heatmaps showing results of three methods (rows) for all cell types (columns).* Each point is colored according to the log2 fold-change, and point size denotes whether the FDR-adjusted p-value is below the 10% level. **(a)** Results for comparing lupus versus healthy samples with Asian ancestry. **(b)** Results for comparing lupus versus healthy samples with European ancestry. **(c)** Inference results on the lupus disease × ancestry interaction effect.

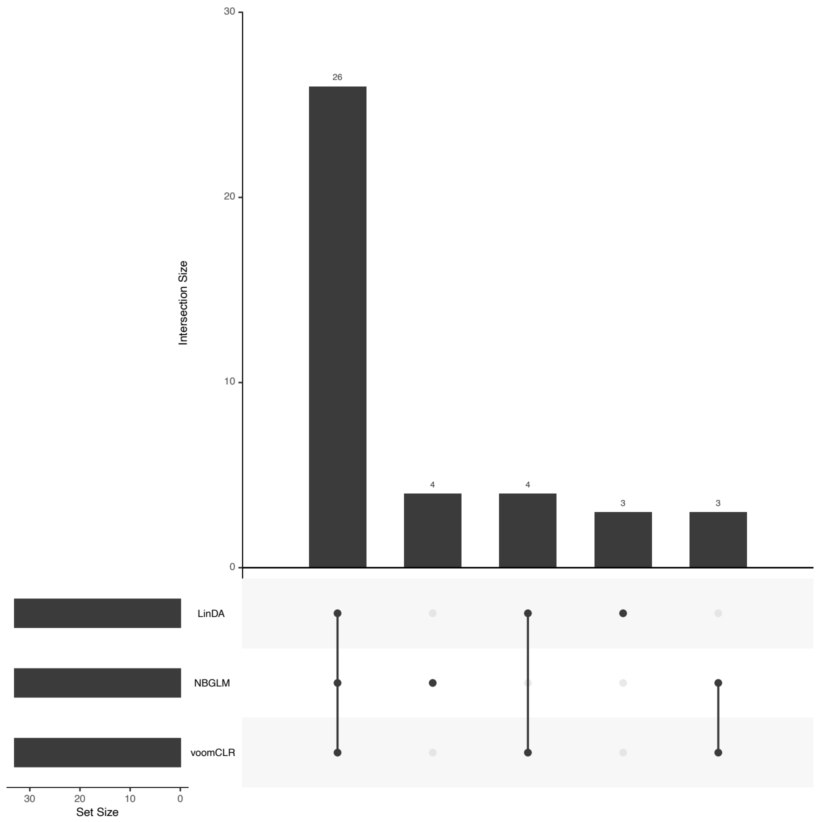


Supplementary Figure S18: *Lupus case study: Upset plot for lupus case study.*

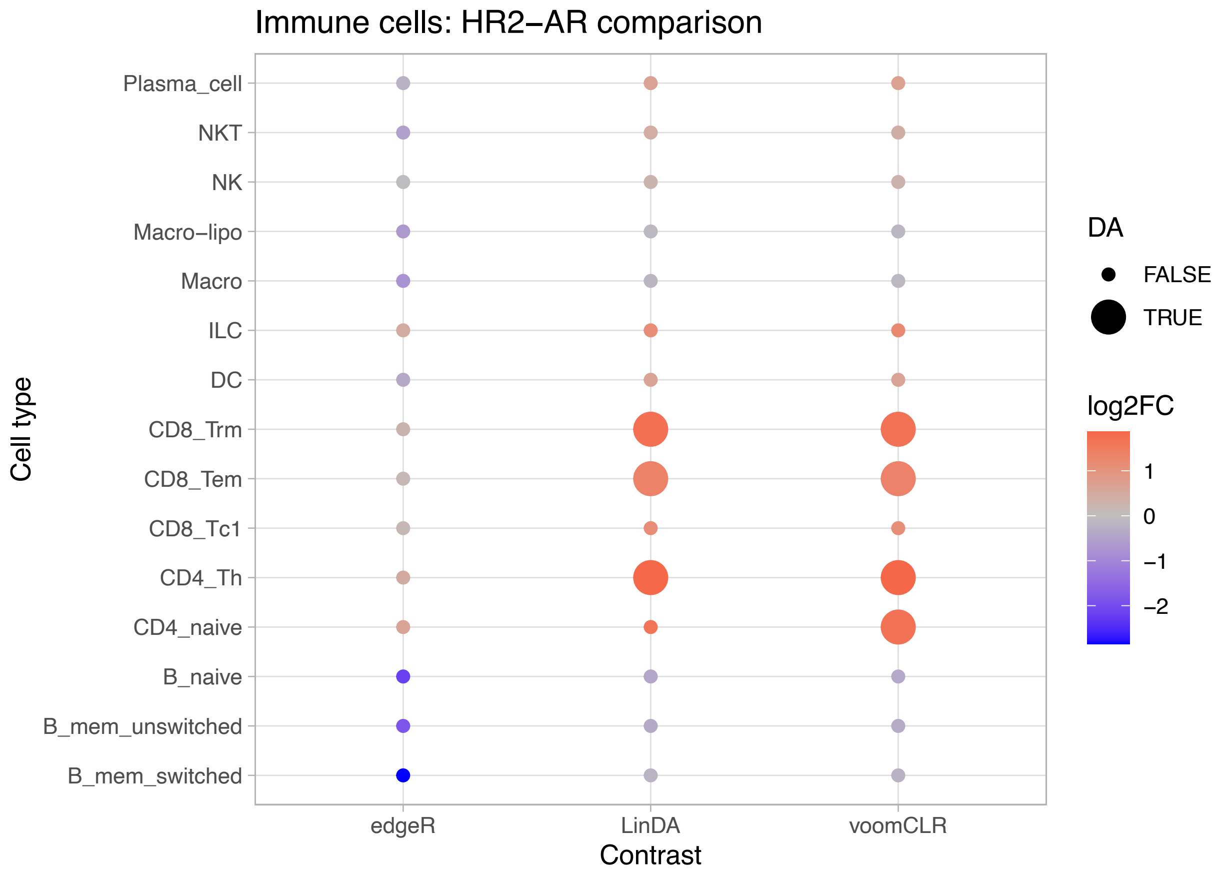


Supplementary Figure S19: *Breast atlas case study: HR2 versus AR donors comparison.* Heatmap showing results of edgeR, LinDA and voomCLR without bootstrapping, for all cell types (y-axis) in the comparison of HR-BR2 versus AR donors. Each point is colored according to the log2 fold-change (positive is higher abundance in HR-BR2 donors), and point size denotes whether the FDR-adjusted p-value is below the 5% level.

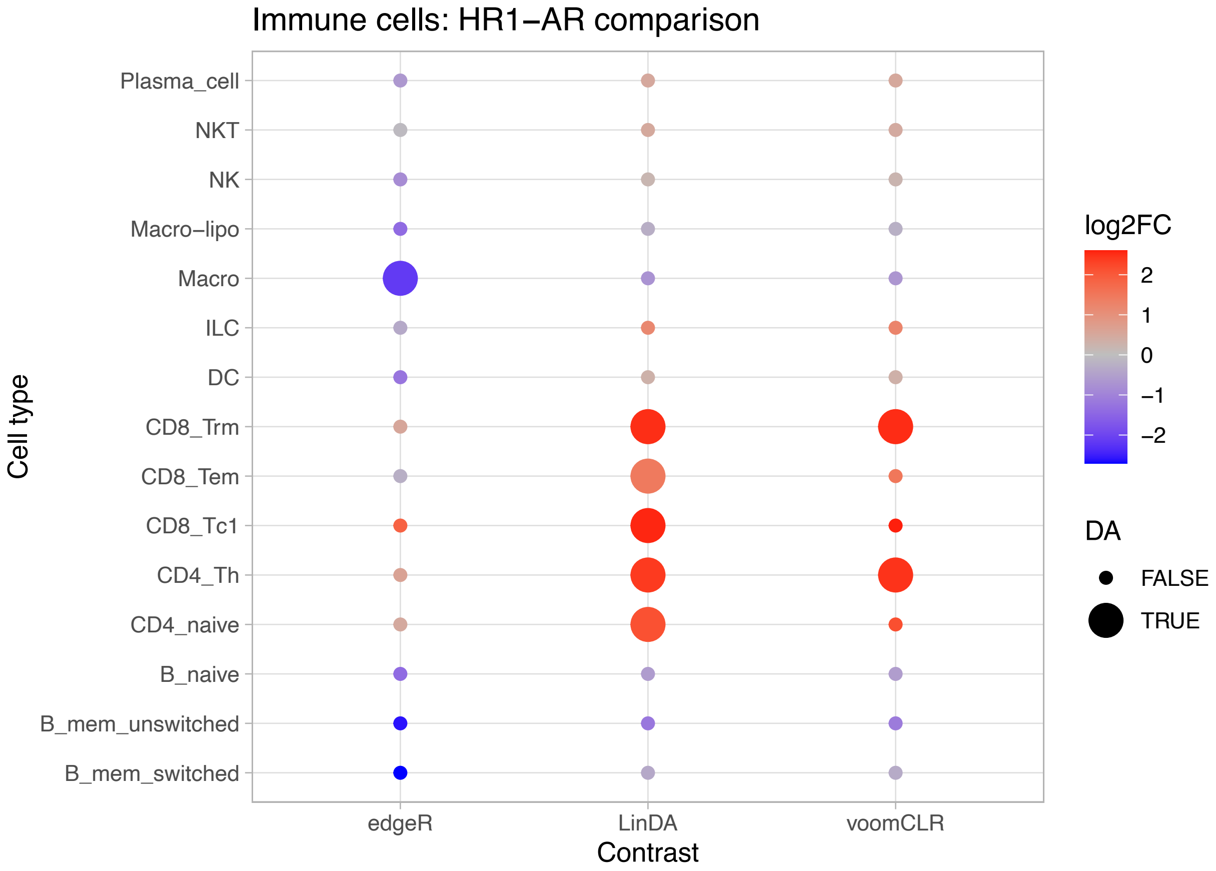


Supplementary Figure S20: *Breast atlas case study: HR1 versus AR donors comparison, using the parametric bootstrap for voomCLR.* Heatmap showing results of edgeR, LinDA and voomCLR with parametric bootstrapping, for all cell types (y-axis) in the comparison of HR-BR1 versus AR donors. Each point is colored according to the log2 fold-change (positive is higher abundance in HR-BR1 donors), and point size denotes whether the FDR-adjusted p-value is below the 5% level.

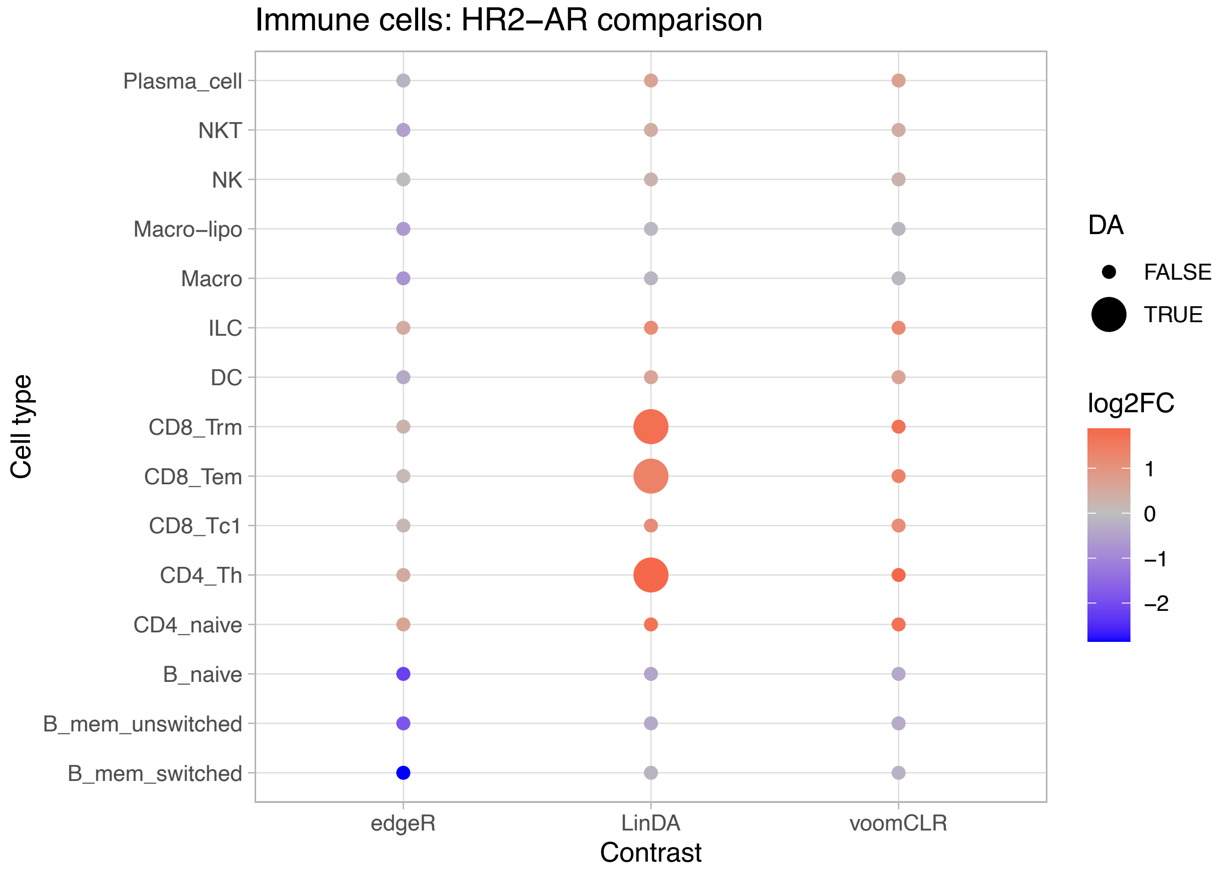


Supplementary Figure S21: *Breast atlas case study: HR2 versus AR donors comparison, using the parametric bootstrap for voomCLR.* Heatmap showing results of edgeR, LinDA and voomCLR with parametric bootstrapping, for all cell types (y-axis) in the comparison of HR-BR2 versus AR donors. Each point is colored according to the log2 fold-change (positive is higher abundance in HR-BR2 donors), and point size denotes whether the FDR-adjusted p-value is below the 5% level.
